# Mutations in core Ccr4-NOT subunits manifest links to decapping rather than deadenylation in *Saccharomyces cerevisiae*

**DOI:** 10.64898/2026.11.19.767818

**Authors:** Agnieszka Tudek, Agnieszka Czarnocka-Cieciura, Rafał Tomecki, Krzysztof Goryca, Aneta Jurkiewicz, Paweł S. Krawczyk, Seweryn Mroczek, Astha Parikh, Matti Turtola, Andrzej Dziembowski

## Abstract

The mRNA polyadenosine tail (pA-tail) is gradually shortened in the cytoplasm by two main deadenylase complexes, Ccr4-NOT and Pan2/3, eventually leading to decapping and decay. The yeast Ccr4-NOT complex contains two deadenylases, Ccr4 and Pop2, and seven non-catalytic subunits. The functions of the latter are not yet fully understood. Here, we investigate the role of Ccr4-NOT in RNA metabolism in budding yeast. We show that steady-state mRNA pA-tail lengths increase with growth temperature, with this effect enhanced in *ccr4Δ* and *pop2Δ* cells, indicating a role for Ccr4-NOT in temperature-dependent mRNA regulation. Consistent with a functional divergence between species, we confirm that Ccr4 is the predominant deadenylase in yeast, whereas the Pop2 orthologs CNOT7/8 are the major deadenylases in human cells. Using Illumina and direct RNA sequencing, we characterize yeast strains defective in Ccr4-NOT, Pan2/3, or mRNA decapping. Unexpectedly, we show that loss of *NOT1*, *NOT2*, *NOT4* or *POP2* causes transcriptome and pA-tail changes resembling those of the decapping-factor mutants *dhh1Δ* and *lsm1Δ*, rather than of the principal deadenylase mutant *ccr4Δ*, and induces a mild stress response. Loss of Not2 or Not4 prolongs the half-lives of several mRNAs and causes growth defects comparable to decapping co-factor mutants. The mRNAs stabilized by loss of Not1-2-4 are distinct from those preferentially deadenylated by Ccr4-Pop2, suggesting different substrate targeting by Ccr4-NOT for non-overlapping functions in deadenylation and decapping.

## INTRODUCTION

The mRNA 3’end polyadenosine tail (pA-tail) is added co-transcriptionally to promote nuclear stability and export to the cytoplasm. Accepted models posit that the pA-tail is gradually shortened during the cytoplasmic mRNA life, and that substantial deadenylation then triggers Dcp2-mediated decapping and degradation from the 5’ end by Xrn1.

The mRNA pA-tails are deadenylated in the cytoplasm by two conserved complexes, Pan2/3 and Ccr4-NOT. Pan2/3 is composed of a single deadenylase, Pan2, and a dimer of Pan3 subunits, which provide the capacity to bind to pA-tails and the polyA-binding protein (Schäfer et al., 2014; Wolf et al., 2014). Ccr4-NOT is a multisubunit complex, which in yeast *Saccharomyces cerevisiae* contains non-catalytic subunits Not1-2-3-4-5, Caf40 and Caf130 along with two main deadenylases, coined Pop2 (Caf1) and Ccr4 (Basquin et al., 2012; Xu et al., 2014). The deadenylases have human orthologs occurring in two isoforms designated CNOT7/8 or CNOT6/6L, respectively. The Ccr4-NOT complex is built around a large scaffolding L-shaped subunit called Not1 in yeast (CNOT1 in humans). Ccr4 and Pop2 bind to one another, and Pop2 directly binds to the central Not1 MIF4G domain near the hinge, thereby connecting Ccr4 to the Not1 scaffold (Basquin et al., 2012; Ukleja et al., 2016; Nasertorabi et al., 2011). The Not2-Not3-Not5 assembly binds at the C-terminus of Not1 (Bhaskar et al., 2013). Deletion of this region of Not1 is lethal (Maillet et al., 2000; Basquin et al., 2012), whereas mutations affecting the interaction of Not2 with Not1 and Not5 confer slow growth at high temperatures (Bhaskar et al., 2013). The Not4 E3 RING ubiquitin ligase associates with Not1 adjacent to the Not2-3-4 module, whereas Caf40 binds the central region of Not1. Caf130 is a yeast-specific subunit associated with the Not1 scaffold (Ukleja et al., 2016).

The non-catalytic core subunits have been assigned several functions in nuclear transcription and cytoplasmic mRNA metabolism. We will only discuss the cytoplasmic functions, as they are most relevant to this study. The Ccr4-NOT complex is recruited to polysomes and the non-catalytic subunits have been shown to play roles in translation regulation. Not4 is required for suppression of translation of aberrant mRNAs, and its loss increases the levels of aggregated proteins, particularly ribosomal subunits. Concomitantly, loss of Not4 reduces the levels of polysomes, and incorporation of Ccr4-NOT into polysomes (Presissler et al., 2015; Panasenko and Collart, 2012). Not5 was also further suggested to monitor mRNA codon optimality contributing to initiation of decay of mRNA with reduced translational efficiency due to problems with decoding (Buschauer et al., 2020). The human orthologs of Caf40, CNOT9, play roles in miRNA-mediated mRNA decay (Chen et al., 2014; Mathys et al., 2014). In budding yeast, RNAi is absent, and the role of Caf40 remains unclear. Caf130 provides a connection between Ccr4–NOT and the ribosome-associated nascent polypeptide-associated complex (NAC), which was shown to be important for translation, folding and ribosomal incorporation of Rpl4 (Schilke et al., 2024). On the other hand, Ccr4-NOT has been shown to partake in deadenylation-independent decapping, which was mediated by the Not2, Not4 and Not5 subunits. This activity was reported for *EDC1* mRNA, because the transcript is resistant to deadenylation (Mühlrad and Parker, 2005). Whether that could be a more general rule remains unclear. Other atudies have also shown physical links between Ccr4-NOT and various decapping co-factors (Maillet et al., 2001; Ozgur et al., 2010; Ukleja et al., 2016). Structural data obtained for the *Schizosaccharomyces pombe* Ccr4-NOT suggested that such physical interaction with decapping co-factors could lead to functional coupling of deadenylation to decapping within the same complex (Ukleja et al., 2016).

Here, we examine the role of budding yeast Ccr4-NOT in adaptation to various temperature growth conditions and assess the function of its core non-catalytic subunits. We show that the steady-state mRNA pA-tail lengths increase with growth temperature, and that this effect is exacerbated in *ccr4Δ* and *pop2Δ* strains, indicating that Ccr4-NOT plays a role in cellular adaptation to changing temperature. Furthermore, we confirm a shift in catalytic importance of the two deadenylases in yeast and human cells. In HEK293 cell lines, it is the yeast Pop2 orthologs, CNOT7/8, that predominantly mediate deadenylation, whereas in yeast Ccr4 is the main deadenylase. To address the function of the Ccr4-NOT core subunits, we produced DRS (direct RNA sequencing) and Illumina datasets from yeast strains with impaired Ccr4-NOT, Pan2/3 subunits, along with those with defective process of decapping. We show that while the scaffolding subunit of Ccr4-NOT in humans, CNOT1, is required for deadenylation, depletion of the yeast counterpart Not1, and deletion of other non-catalytic subunits, results in mRNA pA-tail shortening, which is a phenotype akin to the one observed in decapping co-factor mutant *dhh1Δ* and *lsm1Δ*. The mRNAs with the most shortened pA-tails in the Not1-2-4 defective strains did not overlap with the strongest Pop2-Ccr4 targets, which suggests that Ccr4-NOT might exert two different functions on distinct mRNA classes. Further analyses revealed that changes to mRNA abundance in Not1-deficient, *not2Δ*, *not4Δ* and *pop2Δ* were similar to the ones observed in decapping co-factor mutants *dhh1Δ* and *lsm1Δ*. Accordingly, loss of the *NOT2*, *NOT4*, *POP2* genes conferred growth defects comparable to decapping co-factors *dhh1Δ* and *lsm1Δ*, and far more severe than those observed in main deadenylase mutants *ccr4Δ* and *pan2Δ*. Loss of NOT2 or NOT4 extended the half-lives of several mRNAs, suggesting both proteins exert properties of decapping co-factors.

## RESULTS

### mRNA mean pA-tail lengths increase with growth temperature

To assess the influence of growth temperature on pA-tail metabolism using DRS, we profiled the transcriptomes of wild-type cells grown continuously at 18, 30 and 37°C. This showed that within this 19-degree temperature span, mRNA mean pA-tail lengths increased significantly (Fig. 1a *gray series*), producing an increase of 0.263±0.34 adenosines per degree Celsius. Low-abundant mRNAs displayed slightly higher increases in mean pA-tail lengths (0.27±0.36) compared to medium (0.22±0.2) or high (0.196±0.14) abundant transcripts, but overall the changes were similar for various expression groups (Fig. 1b *black dots*). Altogether, this indicated that growth temperature is a major regulator of steady-state pA-tail length, regardless of mRNA abundance level (Supplementary Fig. 1a-d).

**Figure 1.**
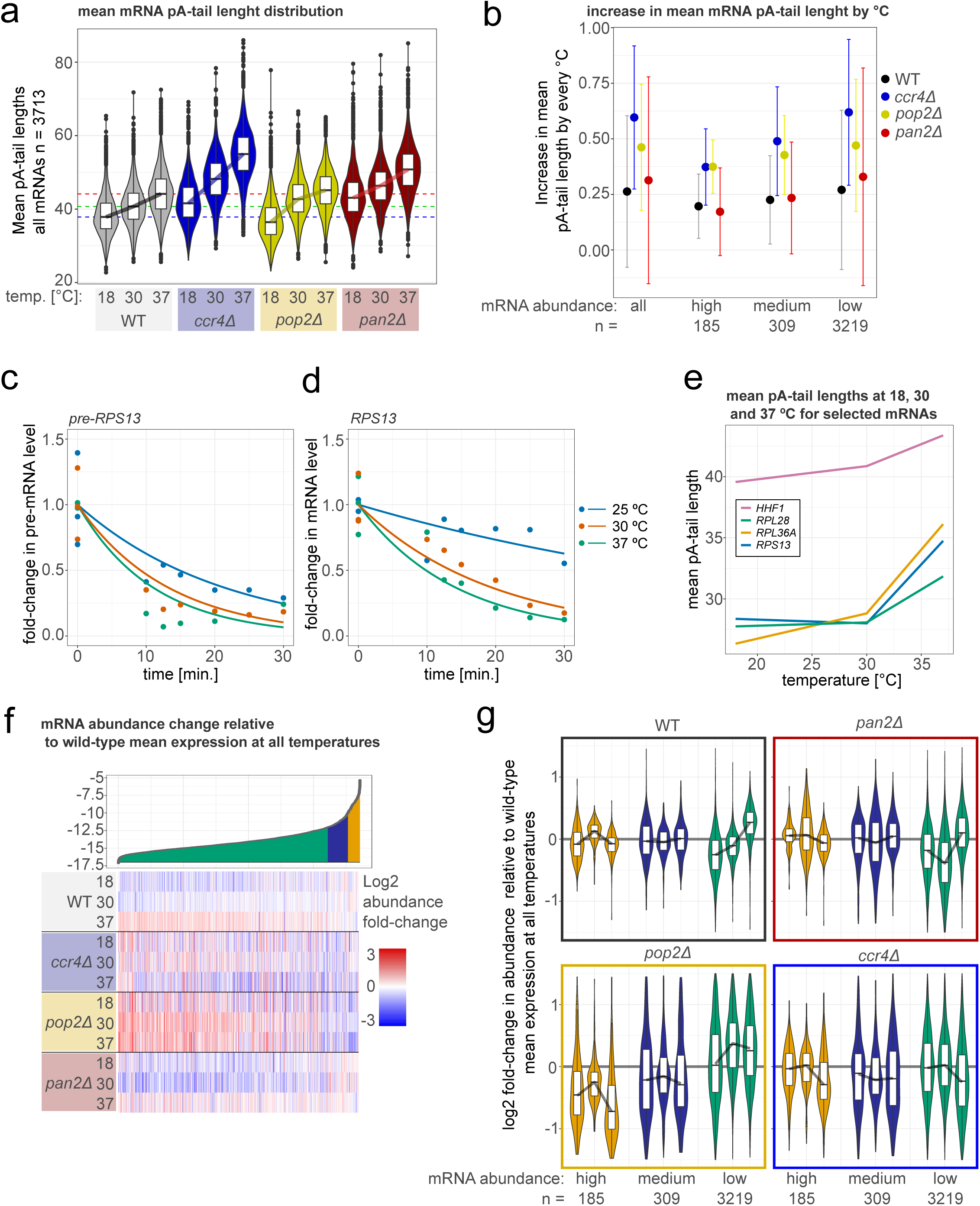
Ccr4-NOT dampens the temperature-dependent increase in pA-tail lengths. **a.** Box and violin plots illustrate the changes to mean mRNA pA-tail lengths for wild-type, *ccr4Δ*, *pop2Δ* and *pan2Δ* strains grown continuously at 18, 30 and 37 °C. **b.** The graph shows the mean increase in individual mRNA pA-tail length calculated for cells grown at 18/30 and 18/37 °C. The error bars represent standard deviations. Values are given for all mRNAs, along with those of high, medium and low abundance per strain. **c-d.** Time-course RT-qPCR analysis of the levels or the *RPS13* mRNA precursor using primers spanning the exon and intron (c), and the *RPS13* mature form (d) in wild-type cells treated with thiolutin at 18, 30 or 37 °C. The single dots represent experimental measurements, while the trend line was inserted using the non-linear fitting protocol in R. **e.** Average pA-tail lengths of *RPS13*, *RPL28*, *RPL36A*, and *HHF1* mRNAs at 18, 30 or 37 °C determined suing DRS. **f.** Heat map shows the mRNA abundance of individual mRNAs in wild-type and *ccr4Δ*, *pop2Δ* and *pan2Δ* strains grown continuously at 18, 30 and 37 °C. The mRNA level changes were normalized to the average expression of each mRNA in wild-type cells grown at 18, 30 or 37 °C. mRNAs are sorter by expression levels, which is demonstrated by the ribbon graph above the heat map. The ribbon is segmented to indicate the position of mRNAs of high, medium and low abundance on the heat map below. **g.** Violin and box plots are an alternative way to present data described in panel f.

We hypothesized that changes in steady-state pA-tail lengths in the wild-type strain could result from variations in mRNA half-lives. As proposed previously by Audebert et al. (2024), more stable mRNAs could allow more time for cytoplasmic deadenylation, resulting in shorter average pA-tails. Conversely, fast decapping and decay would limit the time required for deadenylation, resulting in longer steady-state pA-tails. To test this, we determined the half-lives of several mRNAs at 18, 30 and 37 °C using the transcriptional inhibitor thiolutin (Fig. 1c,d; Supplementary Fig. 1e,f). First, we examined the efficiency of transcription shut-down by assessing the levels of pre-mRNAs through reverse-transcription analysis using oligonucleotides spanning either through the intron-exon junction or over the 3’end cleavage site (Fig. 1c; Supplementary Fig. 1e), as previous works suggested that those intermediates are a good proxy for transcriptional activity (Wu et al., 2022; Schmid et al., 2018). Pre-mRNA levels decreased at a similar rate for cells grown at 30 and 37 °C, whereas transcriptional shut-down was slightly less efficient at 18 °C. The rate of mRNA decay at 18 °C was much slower for transcripts coding for *RPS13*, *RPL28*, *RPL36*, and *HHF1*, even when corrected for less efficient transcriptional shut-down, suggesting overall slower decapping and degradation. In contrast, transcriptional shut-down was of comparable efficiency at 30 and 37 °C, but temperature had no major impact on half-lives at 30 and 37 °C (Fig. 1d; Supplementary Fig. 1f). The selected targets belonged to mRNAs of high expression, and this choice was deliberate to facilitate detection of pre-mRNA forms. However, in this transcript group the increase in mean pA-tail lengths is minor within the window of 18 to 30 °C, but clearly pronounced at 37 compared to 30 °C, and the selected targets followed this pattern (Fig. 1e). Thus, in the case of those mRNAs, half-lives likely decreased in transition from 18 to 30 °C, with no important change in mean pA-tail length. Concomitantly, decay rate remained nearly identical at 30 compared to 37 °C, while average pA-tail lengths noticeably changed. We thus conclude that in the case of temperature growth conditions, alterations in half-lives of mRNA cannot explain changes in mean pA-tail lengths.

### Ccr4-NOT dampens temperature-dependent remodeling of mRNA pA-tails while exerting broader control over transcript abundance

At the moment of nuclear synthesis, the pA-tail is presumed to be of equal size for all mRNAs (Turtola et al., 2021; Tudek et al., 2021), and its length is mainly adjusted by cytoplasmic deadenylases. Therefore, we also profiled the pA-tail lengths of *pan2Δ*, *ccr4Δ* and *pop2Δ* mutants grown at 18, 30 and 37 °C (Fig. 1a,b; Supplementary Fig. 1-d). In all tested conditions, pA-tails in *pan2Δ* were longer than in wild-type, and the mean lengths also increased with temperature. However, the trend of the temperature-dependent increase was identical to the one observed in wild-type cells, regardless of expression group (Fig. 1a,b *red series and dots*). This showed that while Pan2/3 is a major contributor to mRNA deadenylation, it is not responsible for the adaptation to growth temperature.

Mean pA-tails were also longer in *ccr4Δ* compared to control at all respective temperatures. In the case of pA-tails in *pop2Δ,* a more modest elongation was only observed at 30 and 37°C (Fig. 1a; Supplementary Fig. 1-d *blue and yellow series*). In accordance with previous reports, these observations indicated that Ccr4 is the main catalytic subunit of Ccr4-NOT (Czarnocka-Cieciura et al., 2024; Tucker et al., 2002). However, the temperature-dependent increases in pA-tail length in *ccr4Δ* and *pop2Δ* mutants were, respectively, 2.2- and 1.75-fold higher compared to wild-type. Transcripts of low abundance were more robustly affected compared to those of high or medium expression, attaining in *ccr4Δ* and *pop2Δ* an increase of, respectively, 0.62±0.33 and 0.47±0.3 adenosine per degree Celsius (Fig. 1b; Supplementary Fig. 1-d *blue and yellow dots*). This further pointed to a particular role of Ccr4-NOT in cell adaptation to changing growth conditions.

Finally, we asked if pA-tail length changes affected overall mRNA abundance. In wild-type cells, with increasing temperature, transcripts of low abundance experienced an overall increase in mRNA abundance. For mRNAs of high or medium abundance, a clear trend was not observed (Fig. 1f,g). The increase in abundance of mRNAs of low levels was not replicated in deadenylase mutants. However, compared to wild-type, this group of transcripts was up-regulated in *pop2Δ* regardless of growth temperature. This was associated with a concomitant down-regulation of highly abundant mRNAs (Fig. 1f,g).

In sum, temperature-dependent pA-tail elongation promotes gradual accumulation of low-abundance mRNAs in wild-type cells, a balance that is altered in deadenylase mutants, particularly in *pop2Δ*.

### Pop2/CNOT7 is required for proliferation across species despite divergent contributions to deadenylation

Within the Ccr4-NOT complex, the Ccr4 enzyme catalyzes most deadenylation, while Pop2 contributes primarily to the regulation of mRNA abundance (Fig. 1a,f,g; Czarnocka-Cieciura et al., 2024; Tucker et al., 2002). Together, Ccr4 and Pan2 mediate nearly all cellular mRNA deadenylation (Czarnocka-Cieciura et al., 2024), suggesting that Pop2 makes the least significant contribution to deadenylation. Nevertheless, its deletion causes the most severe growth defect among the Ccr4-NOT subunits examined (Fig. 2a). Severely impaired growth was also observed in a Not1 point mutant (F938E/P943Y) defective in its interaction with Pop2 (Fig. 2b; Basquin et al., 2012). Loss of the Ccr4-NOT scaffolding subunit Not1 is lethal (Fig. 2c,d). Together, these observations suggest that the Ccr4-NOT scaffold, through its interaction with Pop2, performs a cellular function that is more essential than deadenylation alone. Notably, among the Ccr4-NOT subunits, loss of Not2 and Not4 were the only other perturbations associated with severely reduced proliferation (Fig. 2a).

**Figure 2.**
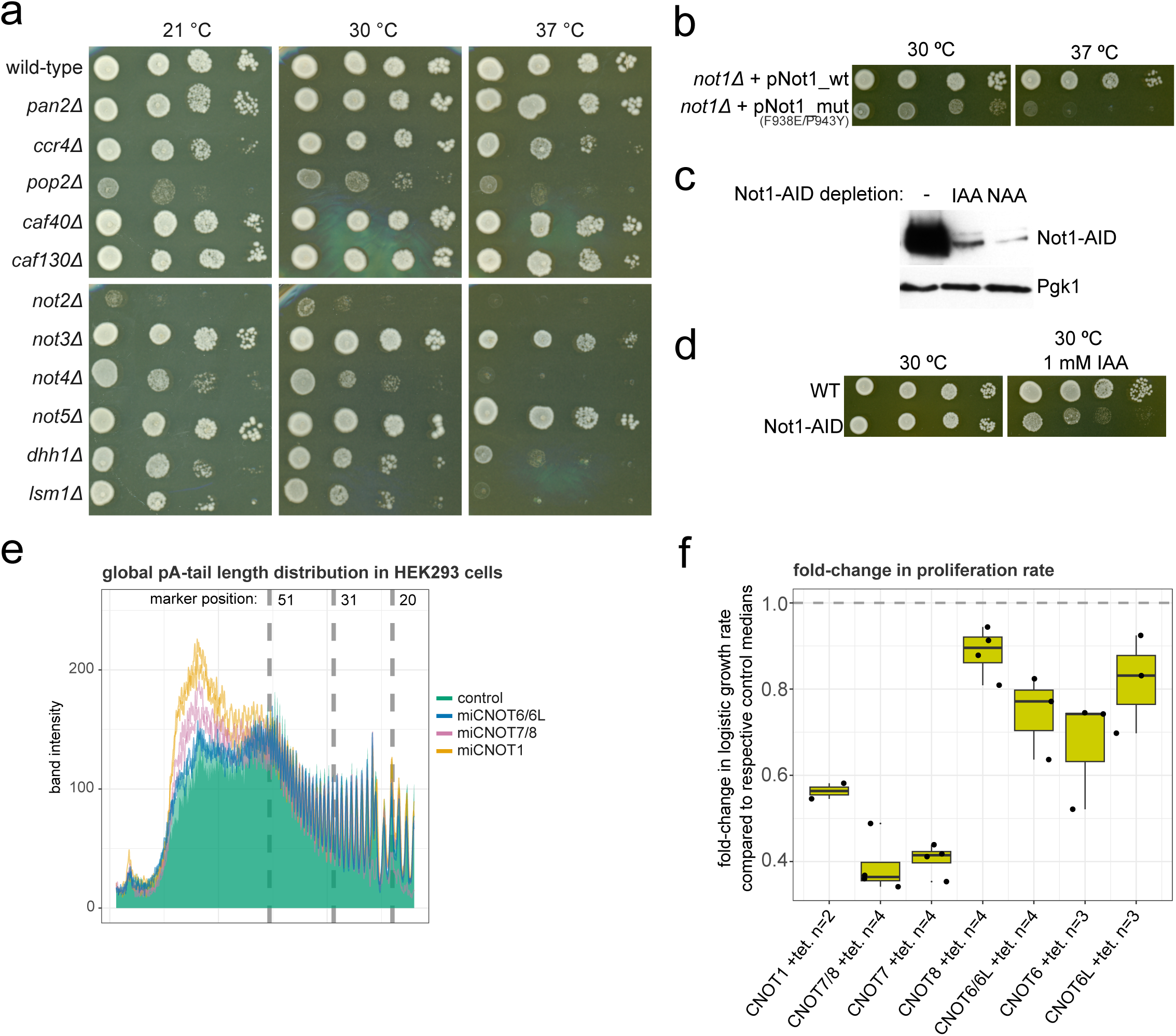
Particular Ccr4-NOT subunits retain their essential character, but diverge in deadenylation roles in yeast compared to humans. **a.** Serial dilution spot test assessing the growth of wild-type W303 yeast or strains with deleted Ccr4-NOT, Pan2/3 subunits, along with *dhh1Δ* and *lsm1Δ*. **b.** Serial dilution spot test assessing the growth of *not1Δ* cells expressing either a wild-type Not1 protein or a mutated version (F938E/P943Y) incapable of interacting with Pop2. **c.** Western blot analysis of the efficiency of Not1 protein depletion using the AID system. Pgk1 protein levels are assessed as a loading control. **d.** Serial dilution spot test assessing the growth of Not1-AID cells in control conditions or in the presence of 1 mM auxin sodium salt. **e.** Graph shows that quantification of the autoradiogram assessing pA-tail lengths in control HEK293 cells compared to cells knocked-down for CNOT7/8, CNOT6/6L or CNOT1 using microRNAs. The source image is shown in Supplementary Fig. 2f. **f.** Box plot showing the fold-change in logistic growth rates in HEK293 cells knocked down for CNOT1, CNOT6, CNOT6L, CNOT7, CNOT8, CNOT6/6L or CNOT7/8. Underlying individual biological replicates are shown as dots.

Previous studies have positioned the human Pop2 orthologs CNOT7 and CNOT8 as the principal catalytic subunits of Ccr4-NOT and have shown that the scaffolding Ccr4-NOT subunit CNOT1 is required for efficient deadenylation (Krawczyk et al., 2025; Mostafa et al., 2020). To further ascertain these findings, we depleted CNOT1, CNOT7, CNOT8, CNOT6 and CNOT6L from HEK293 cells using microRNAs (Supplementary Fig. 2a,b,c,d,e). We then isolated the polyadenylated fraction and assessed pA-tail lengths following RNase A/T1 digestion (Fig. 2e; Supplementary Fig. 2f). Depletion of CNOT1 resulted in an accumulation of pA-tails longer than 50 nt. This effect was weaker, but still significant, following depletion of CNOT7/8, whereas depletion of CNOT6/6L resulted in pA-tail lengths comparable to those of control cells. These results recapitulate previous findings that identify human Pop2 orthologs as the main deadenylase (Mostafa et al., 2020; Yi et al., 2018).

We next assessed the proliferation of HEK293 cells depleted of CNOT proteins. To quantitatively compare experimental confluence curves were modeled growth using a logistic function (Fig. 2f; Supplementary Fig. 2g), as previously described in Tomecki et al. (2026). Comparison of logistic growth rates showed that CNOT1 depletion severely impaired growth. Depletion of CNOT7 alone impaired proliferation, while combined knock-down of CNOT7 and CNOT8 produced only a modest additional effect. Knock-down of either or both CNOT6 or CNOT6L had a modest effect on growth. We conclude that in human HEK293 cells, the scaffolding CNOT1 subunit and Pop2 ortholog CNOT7 are predominantly required for proliferation.

Taken together, these observations indicate that the Ccr4-NOT core scaffolding subunit Not1/CNOT1 and Pop2/CNOT7 are required to sustain efficient proliferation in both budding yeast and human cells. However, their contributions to deadenylation differ between the two systems. In human cells, Pop2/CNOT7 contributes substantially to deadenylation, whereas in yeast this function is predominantly performed by Ccr4. Thus, the contribution of individual Ccr4-NOT catalytic subunits to bulk deadenylation does not directly predict their importance for cellular proliferation and appears to have diverged between yeast and humans. These findings suggest that the essential functions of Ccr4-NOT extend beyond its role in bulk mRNA deadenylation and highlight budding yeast as a convenient system for identifying the essential functions of the core Ccr4-NOT complex that are uncoupled from its major deadenylase activity.

### The core of the Ccr4-NOT complex is dispensable for deadenylation and loss of its subunits leads to pA-tail shortening akin to phenotypes observed in decapping mutants

To evaluate the role of the yeast Ccr4-NOT non-catalytic subunits in deadenylation, we produced a panel of DRS datasets (Fig. 3a). Since *NOT1* is an essential gene, we used the auxin inducible degron system to remove the protein (Nishimura et al., 2009; Fig. 2c,d), and we also sequenced the *not1Δ* strain in which the gene was complemented by expression of wild-type Not1 protein or the mutant incapable of binding to Pop2 (F938E/P943Y; Fig. 2b; Basquin et al., 2012). Depletion of Not1 using the AID tag shortened mRNA pA-tails by around 3.3 adenosines on average (Fig. 3a). This phenotype contrasts with previous observations in human HEK293 cell lines where depletion of CNOT1 resulted in pA-tails elongation by 12.5 adenosines (Fig. 3b; Fig. 2e; Krawczyk et al., 2025). In addition, the Not1 mutant incapable of binding to Pop2 also displayed pA-tail shortening by 3.8 adenosines. Altogether, this supports the notion that the scaffolding subunit of Ccr4-Not does not perform the same function in yeast compared to humans.

**Figure 3.**
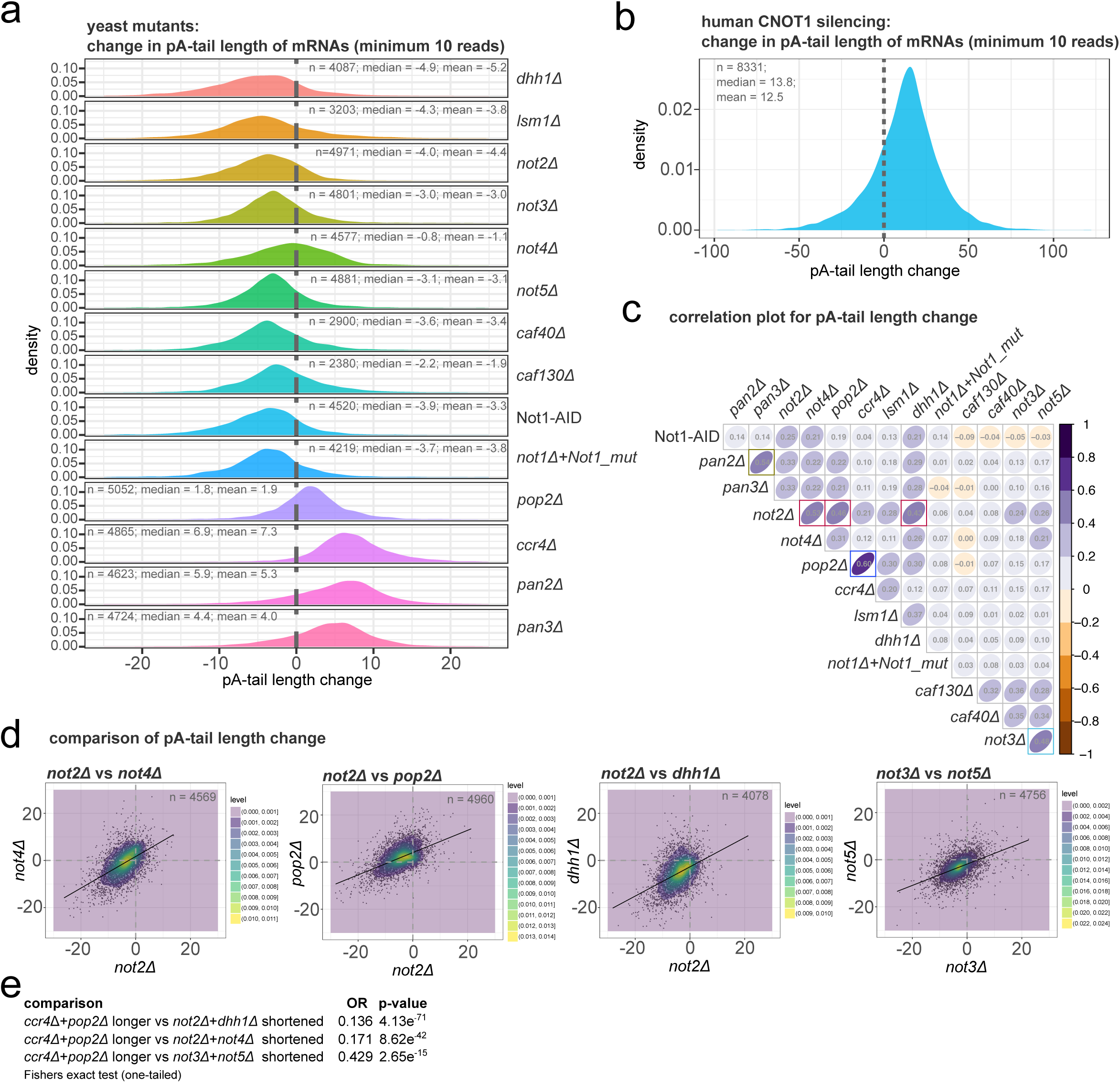
Loss of yeast core Ccr4-NOT subunits results in mRNA pA-tail length shortening akin to decapping mutants. **a.** Stacked density plots show the distribution of pA-tail length changes in indicated yeast Ccr4-NOT, Pan2/3 or decapping co-factor mutants compared to wild-type. The mean, median and number of mRNAs observed are indicated on the panel. **b.** Stacked density plots show the distribution of pA-tail length changes in HEK293 cells knocked down for CNOT1 compared to non-depleted control. The mean, median and number of mRNAs observed are indicated on the panel. **c.** Matrix showing mean pA-tail length Spearman ρ correlation coefficients calculated pairwise for yeast strains impaired in *NOT1-2-3-4-5*, *CAF40130*, *CCR4*, *POP2*, *PAN2/3*, *DHH1* and *LSM1* function. The number of pairwise observations is listed in Supplementary Fig. 3a. **d.** Scatter plots compare mRNA pA-tail length changes relative to wild-type between *not2Δ*-*not4Δ*, *not2Δ*-*pop2Δ*, *not2Δ*-*dhh1Δ*, and *not3Δ-not5Δ*. The number of pairwise observations is indicated on each graph. A two-dimensional density plot is overlaid on top of the scatter, and a linear regression line is drawn through the data. The Spearman ρ correlation coefficients are shown on panel c. **e.** The table lists results of the Fisher exact test and odds ratio (OR) for groups of mRNAs with elongated pA-tails in *ccr4Δ*-*pop2Δ* compared to groups of transcripts with shortened pA-tails in *not2Δ*-*not4Δ*, *not2Δ*-*dhh1Δ*, and *not3Δ-not5Δ*.

The DRS dataset showed that mRNA pA-tails were shortened in all non-catalytic Ccr4-Not deletion mutants (Fig. 3a). The strongest pA-tail shortening, by 4.4 adenosines, was observed in the *not2Δ* strain, and the most modest in *not4Δ* (1.1 As). The *not3Δ*, *not5Δ*, *caf40Δ* and *caf130Δ* mutants showed average pA-tail shortening by 2.2 to 3.3 adenosines. For comparison, loss of *CCR4* or *POP2* elongated mRNA pA-tails by 7.3 or 1.9 adenosines on average, respectively (Czarnocka-Cieciura et al., 2024; Tudek et al., 2021). We previously observed that depletion of the decapping factor Dcp2 or exonuclease the Xrn1 resulted in length restriction of mRNA pA-tails (Czarnocka-Cieciura et al., 2024). To strengthen this observation, we sequenced transcriptomes of the *dhh1Δ* and *lsm1Δ* strains, devoid of decapping co-factors. In those strains, we again observed pA-tail length shortening by 5.2 and 3.8 adenosines (Fig. 3a). We conclude that mutations in the Ccr4-NOT scaffolding and non-catalytic subunits result in global shortening of pA-tails, which is a phenotype also observed in decapping factors.

Next, we compared the changes in individual mRNA pA-tail length between strains (Fig. 3c,d and Supplementary Fig. 3a,b). As expected for enzymes partaking in the same complex, this revealed robust similarities between the *ccr4Δ* and *pop2Δ* strains (Spearman ρ=0.6), and also separately between *pan2Δ* and *pan3Δ* (ρ=0.54). Loss of Pan2/3 or Ccr4 did not produce any marked correlation, in accordance with previous notions that both deadenylases display their own substrate preferences (Tudek et al., 2021). Our analysis also revealed two other similarity clusters, between *not3Δ* and *not5Δ* (ρ=0.48), and between *not2Δ* and three other strains, *not4Δ, pop2Δ* or *dhh1Δ* (ρ=0.53, 0.46, 0.43, respectively).

We hypothesized whether shortened pA-tails observed in *not* mutants could correspond to excessively deadenylated targets of Ccr4-Pop2 deadenylases. We reasoned that ongoing Ccr4-Pop2-mediated deadenylation could trigger decapping, but in the absence of the core Not subunits, decapping would be delayed, resulting in shorter tails. However, cross comparison of mRNA groups which have elongated pA-tails in *ccr4Δ* or *pop2Δ* mutants, to those that have shortened pA-tails in *not2Δ*-*not4Δ* or *not2Δ*-*dhh1Δ* mutants revealed an unexpectedly low overlap (Fisher exact test OR ratio 0.171 or 0.136; with high significance), indicating that the groups are in fact distinct. The OR ratio was substantially less robust for the *not3Δ*-*not5Δ* group (0.429) but still significant (Fig. 3e). Further gene ontology analysis indicated that each set of targets grouped mRNAs functionally relevant in different biological processes. Targets of *ccr4Δ* and *pop2Δ* grouped 901 transcripts relevant to mitosis regulation, DNA repair, and cell-cycle checkpoint (Supplementary Figure 3c). The *not3Δ-* and *not5Δ*-dependent mRNAs (n = 779) were mainly involved in rRNA, sn/snoRNA, and tRNA maturation, polyadenylation-dependent mRNA catabolic processes, and DNA repair (Supplementary Figure 3d). In the *not2Δ* similarity cluster, the largest number of gene ontology groups were formed with *dhh1Δ* (n = 1093), spanning groups of mRNAs involved in RNA polymerase II pre-initiation complex assembly, RNA capping, cytosolic transport and protein targeting to membrane, trehalose and glycogen metabolism, proteasome function and ERAD pathway, and mitochondrial respiratory function (Supplementary Figure 3e). The *not2Δ*-*pop2Δ* (n = 242) and *not2Δ*-*not4Δ* (n = 734) groups produced few gene ontology groups, sharing the trehalose biosynthetic process with the *not2Δ*-*dhh1Δ* cluster (Supplementary Figure 3f,g).

Given this data, we conclude that loss of core Ccr4–NOT proteins Not2-3-4-5 and decapping co-factor Dhh1 results in pA-tail shortening of transcript populations that are distinct from the mRNAs showing increased pA-tail length when Ccr4–Pop2-dependent deadenylation is perturbed. This possibly indicates that they represent a separate target set for the Ccr4-NOT complex that is not strongly dependent on deadenylation by Ccr4-Pop2.

### pA-tail shortening is linked to an increase in mRNA abundance in Not2, Not4 and Not1-impaired strains

Increase in mRNA abundance is one of the simplest indicators of impaired decapping and degradation. Therefore, we analyzed the changes to mRNA abundance in our DRS datasets (Figure 4a). Out of roughly 5000 protein-coding transcripts, only 2133 were detected in all sequenced strains, clearly limiting our insight. However, we observed an up-regulation of mRNAs of low expression in Ccr4-NOT-, Dhh1- and Lsm1-impaired strains, which was not observed in *PAN2/3* deletions. This occurred with concomitant down-regulation of mRNAs of high abundance. This phenotype was most marked in *not2Δ* and *pop2Δ* mutants (Figure 4a, left panel). Next, we performed hierarchical clustering of mutants based on Spearman correlation of genome-wide mRNA abundance changes, using average linkage (Figure 4a, right panel). This grouped *not2Δ* and *not4Δ* in one cluster, which was linked to a larger group composed of *dhh1Δ*, *lsm1Δ*, *pop2Δ*, *ccr4Δ*, and both the Not1 depletion and the Not1 mutant incapable of binding to Pop2.

**Figure 4.**
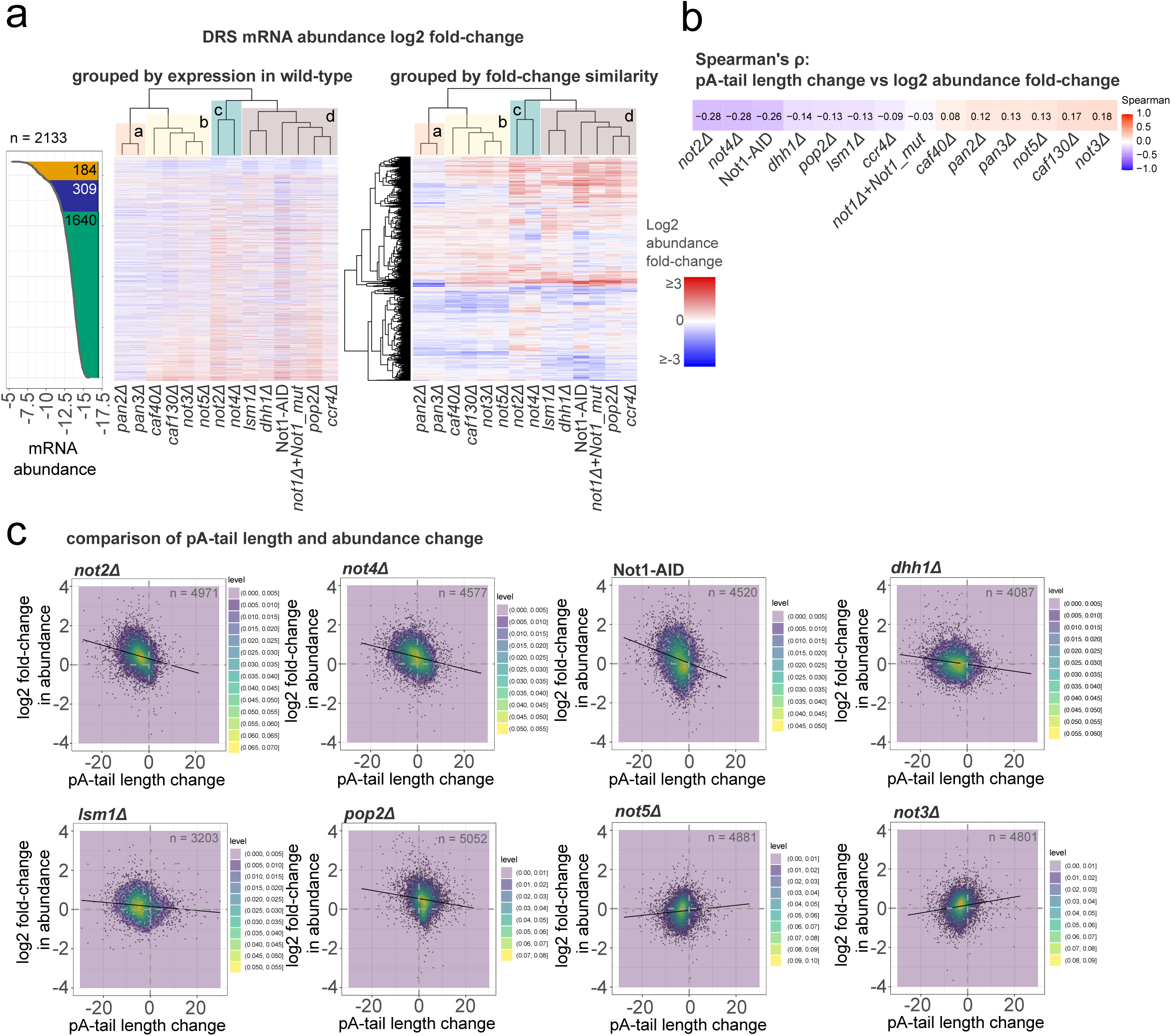
Loss of Not1-2-4 and decapping co-factors results in simultaneous pA-tail shortening and mRNA up-regulation. **a.** Heat maps assessing changes to mRNA abundance in DRS datasets derived from strains impaired for Dhh1, Lsm1, and Ccr4-NOT or Pan2/3 subunits relative to wild-type. The heat map on the left sorts mRNAs by expression level, and to indicate mRNAs of high, medium and low expression, a neighboring ribbon plot is presented. The heat map on the right groups mRNAs by similarity in expression change between strains. Strains are also grouped by similarity using the hclust protocol in R. **b.** The colored table lists the Spearman ρ coefficients in mRNA mean pA-tail length change and log2 abundance fold-change for the DRS datasets. The number of observations is equal to the number given in Fig. 3a. **c.** Scatter plots show pA-tail length and log2 abundance changes in mutant strains (*not2Δ*, *not4Δ*, *dhh1Δ*, *lsm1Δ*, *pop2Δ*, *not5Δ*, *not3Δ*) or in cells depleted of Not1 using the AID system compared to wild-type. The number of mRNAs examined is given as ‘n =’. A linear regression line is drawn; the Spearman ρ coefficients are listed in panel b.

Next, we correlated the changes in pA-tail length and abundance (Figure 4b). Negative Spearman ρ coefficients were identified for *not2Δ*, *not4Δ*, *pop2Δ*, *dhh1Δ*, *lsm1Δ*, and Not1-depleted strains. Closer inspection of the data (Figure 4c; Supplementary Fig. 4a) indeed revealed a clear relationship between pA-tail length restriction and an increase in mRNA abundance, especially for the *not2Δ*, *not4Δ*, and Not1-depleted strains, and a milder one for *dhh1Δ* and *lsm1Δ* strains. Despite a clear pA-tail length shortening in *not3Δ*-*not5Δ,* we observed no such relationship with mRNA abundance. Together, these results identify the Not2-Not4-Not1 proteins as potential factors coupling mRNAs to decapping in a manner analogous to Dhh1 and Lsm1.

### Differential expression changes in the Not1, Not2, Not4, and Pop2-impaired cells position them in phenotypic vicinity to Dcp2-depleted cells and decapping co-factor mutants *dhh1Δ* and *lsm1Δ*

The mRNA abundance data gathered in the course of DRS dataset preparation were promising, but the moderate read coverage limited the strength of the conclusion. We therefore prepared new samples and subjected them to Illumina sequencing using anchored oligo-dT primed libraries. This substantially increased the read coverage and enabled us to better determine RNA differential expression status, which now covered nearly the entire coding transcriptome (5078 mRNAs; Fig. 5a). The log_2_ mRNA abundance changes in the Illumina dataset mostly correlated well with the DRS data, attaining Spearman ρ values of 0.48 to 0.74 for strains deficient for Not1-2-4, Pop2, Ccr4, Dhh1, and Lsm1. Poor correlations were observed for the rest of the strains, namely *not3Δ*, *not5Δ*, *caf40Δ*, *caf130Δ*, and *pan2Δ*. However, those were strains where, in general, little significant changes in mRNA levels were noted, as reflected by the low median absolute deviation of the log_2_ mRNA abundance fold-change in both the DRS and Illumina datasets (Fig. 4a, 5a,b). Overall, this degree of mRNA abundance variability in the Illumina and DRS datasets showed a good correlation with Spearman ρ – 0.69 (Fig. 5c).

**Figure 5.**
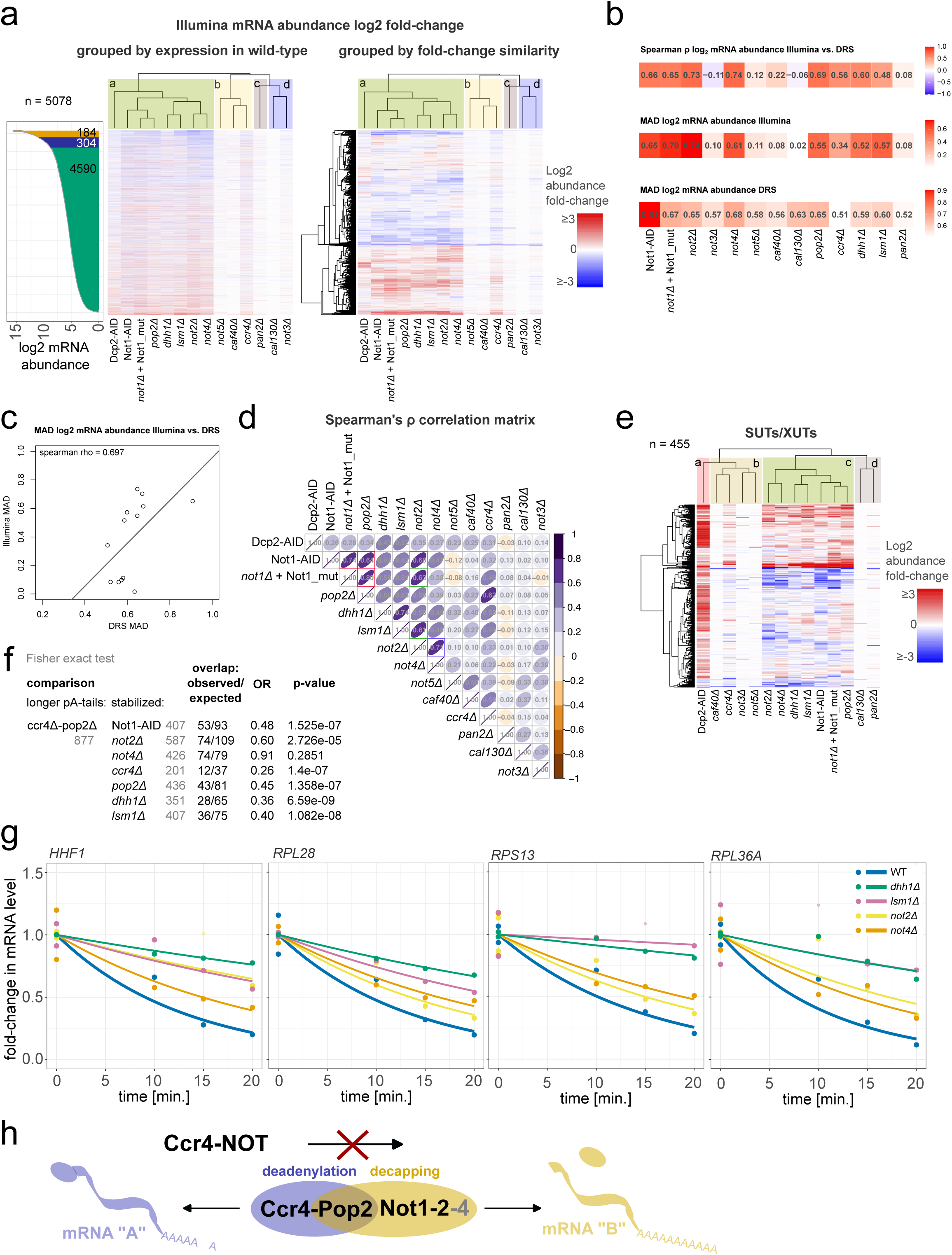
Loss of the Not1-2-4 core Ccr4-NOT subunits and Pop2 manifests phenotypic similarities to deletions of decapping co-factors *DHH1* and *LSM1*. **a.** Heat maps show mRNA abundance changes in oligo-dT-primed Illumina datasets derived from strains impaired for Dcp2, Dhh1, Lsm1, and Ccr4-NOT or Pan2/3 subunits relative to wild-type. The heat map on the left sorts mRNAs by expression level to indicate mRNAs of high, medium and low expression. A neighboring ribbon plot is presented. The heat map on the right groups mRNAs by similarity in expression change between strains. Strains are also grouped by similarity using the Spearman ρ coefficients (see panel d). **b.** Colored tables list from top to bottom the Spearman ρ coefficients of log2 mRNA abundance fold-change relative to wild-type in DRS versus Illumina datasets (top), the median absolute deviation (MAD) in log2 mRNA abundance fold-change relative to wild-type for the Illumina (middle) and DRS datasets (bottom). **c.** The scatter plot compares the MAD values in the DRS and Illumina datasets. A linear regression line is drawn, and the Spearman ρ coefficient, equal to 0.69, is indicated. **d.** Matrix of Spearman ρ coefficients correlating the log2 mRNA abundance fold-changes in various strains examined using the Illumina protocol. The number of observations is 5078 for all assessed strains. **e.** Heat map shows SUT/XUT abundance changes in oligo-dT-primed Illumina datasets derived from strains impaired for Dcp2, Dhh1, Lsm1, and Ccr4-NOT or Pan2/3 subunits relative to wild-type. The heat map is sorted by ncRNA expression similarity, while the strains are grouped by similarity of Spearman ρ coefficients. **f.** The table sums up the results of one-tailed Fisher exact test comparing populations of mRNAs with pA-tails elongated by at least 5 adenosines in *ccr4Δ* and *pop2Δ* depleted cells in DRS datasets to mRNAs stabilized by at least two-fold in the Illumina dataset in Not1-AID, *not2Δ*, *not4Δ, ccr4Δ*, *pop2Δ*, *dhh1Δ* and *lsm1Δ* strains. **g.** RT-qPCR analysis of *HHF1*, *RPL28*, *RPS13*, and *RPL36A* abundance in thiolutin time-course for wild-type, *not2Δ*, *not4Δ*, *dhh1Δ* and *lsm1Δ* strains. The precursor levels are shown in Supplementary Fig. 5d. **h.** Model showing the dual function of the Ccr4-NOT complex. While the Ccr4-Pop2 deadenylases mediate pA-tail shortening, the core subunits Not1, Not2, Not4, along with Pop2, are linked to decapping. The populations of mRNAs targeted by each Ccr4-NOT function are not overlapping.

The DRS dataset was prepared on the polyadenylated fraction, whereas the Illumina sequencing utilized anchored oligo-dT priming. The potential for material loss was therefore greater in the DRS compared to the Illumina dataset. In the case of the *not3Δ*, *not5Δ*, *caf40Δ* and *caf130Δ* strains, where marked changes in pA-tail length were noted in DRS data, oligo dT selection likely translated into more important changes in mRNA abundance, which were not detected in the Illumina dataset (Fig. 4a, 5a). Thus, those Ccr4-NOT subunits influence more strongly the polyadenylation status of mRNAs, rather than their stability.

The Illumina analysis clearly revealed that within the Ccr4-NOT complex it is the Not1 scaffold, Not2, and Not4 that have more impact on the overall mRNA abundance. Curiously, out of the two deadenylases, loss of Pop2 had a more dramatic influence on mRNA abundance compared to Ccr4. This role of Pop2 was clearly dependent on the interaction with the Ccr4-NOT core, as the Not1 mutant incapable of binding to Pop2 also displayed large changes in mRNA abundance (Fig. 5a,b). Overall, changes to mRNA abundance correlated well within the set of strains with depleted Not1 scaffold, *not2Δ*, *not4Δ*, *pop2Δ*, and the decapping co-factors *dhh1Δ* and *lsm1Δ* (Fig. 5a,d), whereas strains deficient for *CCR4* or *DCP2* displayed a more distant similarity. Comparably to the DRS dataset, this strain cluster produces a marked tendency for up-regulation of mRNAs of low expression, and concomitant down-regulation of abundant mRNAs (Fig. 4a, 5a).

The Illumina dataset sequencing depth allowed for more detailed analysis of the status of non-coding RNAs (Fig. 5e, Supplementary Fig. 5a,b,c). SUTs and XUTs are the ncRNA classes that are most markedly dependent on Dcp2 (van Dijk et al., 2011; Marquardt et al., 2011). Depletion of Dcp2 using the AID system resulted in near-uniform stabilization of XUTs/SUTs. However, loss of the *DHH1* and *LSM1* decapping co-factors only resulted in stabilization of roughly a third of those RNAs, with concomitant down-regulation of the remaining pool. This pattern was reproduced in the Not1-2-4 and Pop2-deficient strains (Fig. 5e). Similar observations were made with the detectable pool of RNAs arising from CUT loci (Supplementary Fig. 5a). Loss of Not1-2-4 or Pop2 also impacted snRNA processing, along with some tRNAs, and *NTS2-1* and *NTS1-2* transcripts, arising from the rRNA locus (Supplementary Fig. 5b,c). In the case of SUTs/XUTs and detectable CUTs, which likely have a limited cytoplasmic fraction, we can presume that Ccr4-NOT acts as a co-factor, similar to Dhh1 and Lsm1 in the decay of those RNAs.

Comparison of DRS pA-tail estimates with Illumina abundance measures showed that a pA-tail length shortening was associated with increased mRNA abundance (Supplementary Fig. 5d), which resonated with the DRS dataset analysis presented in the previous section (Fig. 4c and Supplementary Fig. 4a). This was most pronounced for *not2Δ* (ρ = -0.42), followed by *not4Δ* (-0.37), *lsm1Δ* (-0.35), Not1 depletion (-0.33), and *dhh1Δ* (-0.32), with marked cross-similarities between those strains. Importantly, this association was also observed, however with lesser strength, in *pop2Δ* and *ccr4Δ* mutants (-0.24 for both). Further analyses showed that transcripts that accumulate long pA-tails upon impairment of Ccr4/Pop2-dependent deadenylation are significantly depleted among transcripts stabilized by loss of *ccr4Δ*, *pop2Δ*, Not1-AID, *not2Δ*, *dhh1Δ*, or *lsm1Δ*, as supported by Fisher exact test results (Fig. 5f). This supported the previously proposed notion that the populations targeted for deadenylation or decapping by Ccr4-NOT are largely distinct.

To determine whether Not2 and Not4 could function as decapping co-factors similar to Dhh1 and Lsm1, we asked whether loss of either protein impacted mRNA half-lives. To this end, we performed a thiolutin chase at 30 °C. The levels of nascent mRNAs dropped in each deletion mutant similarly to wild-type (Supplementary Fig. 5e), indicating successful transcriptional shut-down. As expected, loss of *DHH1* and *LSM1* almost entirely blocked the decay of *HHF1*, *RPS13*, *RPL28* and *RPL36A* mRNAs. Deletion of *NOT2* and *NOT4* had a less dramatic effect, but still markedly prolonged the half-lives of all tested transcripts (Fig. 5g).

Together, these results suggest that the Not2-Not4-Not1 proteins may be responsible for coupling decapping and decay in a way analogous to *bona fide* decapping co-factors Dhh1 and Lsm1. However, the mRNAs that are targeted are distinct from the preferred Ccr4-Pop2 deadenylation targets. This could indicate that Ccr4-NOT complex performs two independent functions as a deadenylase and as a decapping co-factor (Fig. 5h).

### Loss of *NOT1-2-4*, *DHH1* and *LSM1* induces a mild stress response

Finally, we asked which groups of mRNAs were commonly stabilized or down-regulated in mutants of core Ccr4-NOT subunits, *not2Δ*, *not4Δ* and Not1-AID depletion, and also in decapping co-factor strains *dhh1Δ* and *lsm1Δ*. This yielded a set of 363 mRNAs stabilized or 157 down-regulated by at least two-fold in at least three out of the five analyzed strains (Fig. 6a). Functionally, each group formed very few gene ontology groups (Supplementary Figure 6a). Up-regulated mRNAs played roles in carbohydrate metabolism, including TCA cycle activity, pentose degradation, UDP-GlcNAc biosynthesis, and regulation of gluconeogenesis. Down-regulated transcripts were linked to amino acid metabolism and translation/ribosome biogenesis.

**Figure 6.**
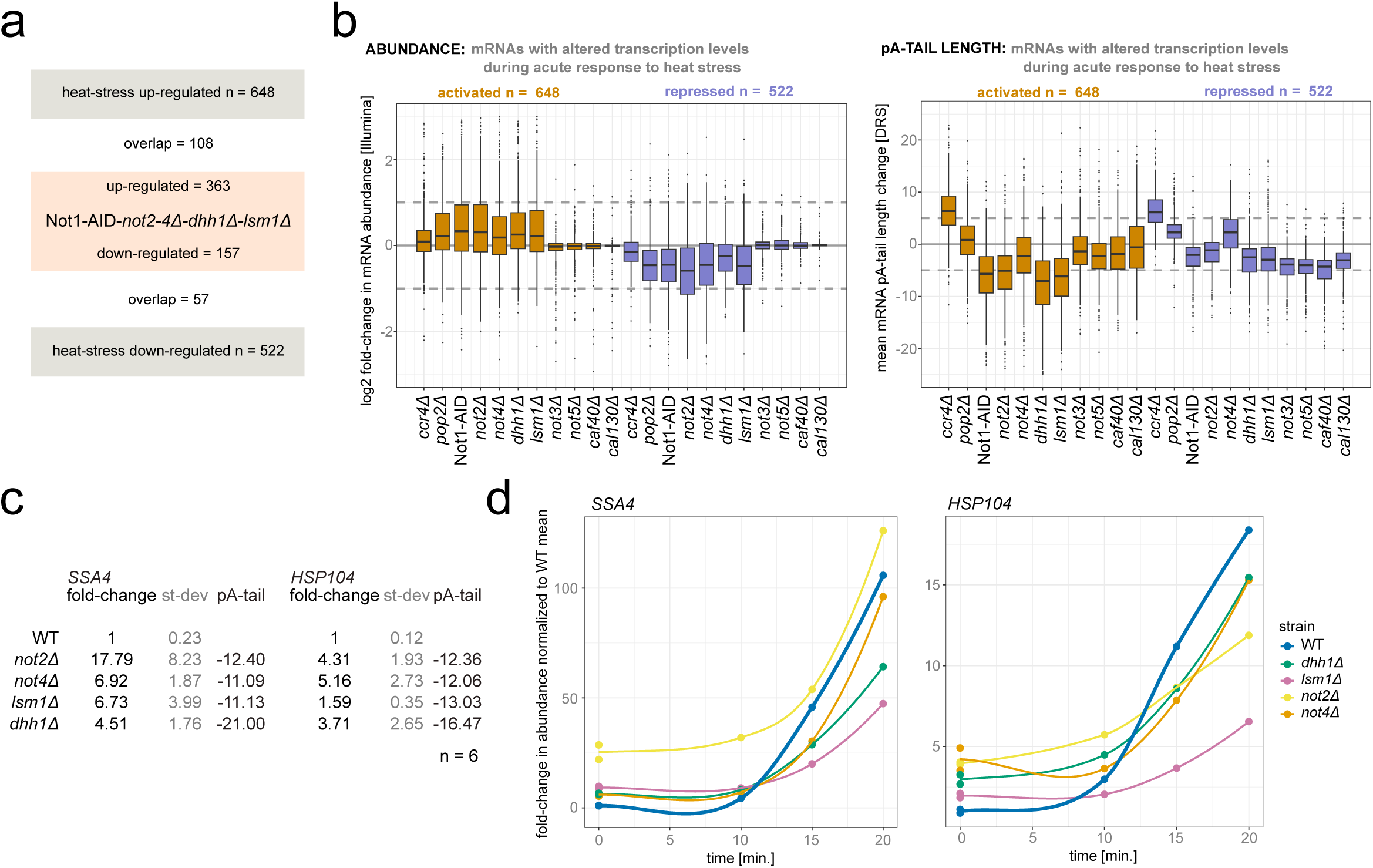
Loss of Not1-2-4 and Dhh1, Lsm1 induces a mild stress response. **a.** Panel examines the number of mRNAs up- or down-regulated in Not1-AID, *not2Δ*, *not4Δ*, *dhh1Δ* or *lsm1Δ*. mRNAs were incorporated if differentially expressed by at least 2-fold in 3 out of 5 strains. **b.** Box plots examine mRNA abundance in the Illumina datasets (left) or mean pA-tail length change in DRS (right) of mRNA previously reported to be transcriptionally silenced or activated during acute response to heat stress by Vinayachandran et al. (2018). **c.** The table lists the SSA4 and HSP104 mRNA fold-change assessed by RT-qPCR along with mean pA-tail length change in DRS for *not2Δ*, *not4Δ*, *dhh1Δ* or *lsm1Δ* strains. **d.** Graphs show up-regulation of *HSP104* and *SSA4* mRNAs by RT-qPCR in wild-type, *not2Δ*, *not4Δ*, *dhh1Δ* or *lsm1Δ* strains exposed to low levels of thiolutin. Trend lines were produced with locally estimated scatterplot smoothing. Single experimental measurements are shown as dots. Data were normalized to the average levels of each mRNA in wild-type cells.

Manual inspection of the gene groups revealed consistent up-regulation of heat-shock-induced mRNAs, *HSP12-26-30-42-78-82-104*, and *SSA4*, and concomitant down-regulation of 24 ribosomal protein-coding transcripts (Supplementary Figure 6a). This is a transcriptional pattern previously reported for the acute response to heat shock (Vinayachandran et al., 2018), during which severe protein misfolding occurs. Notably, Not4 was also reported to be required for the maintenance of cellular proteostasis (Preissler et al., 2015). Initial cross comparison of mRNAs jointly changed in abundance in Not1-AID-*not2Δ-not4Δ-dhh1Δ-lsm1Δ* group revealed that the differential expression of only roughly a third of those mRNAs could be explained by transcriptional changes previously defined for acute response to heat stress (Fig. 6a). However, reverse analysis of the behavior of mRNA groups defined by Vinayachandran et al. (2018) showed that in the Not1-AID-*not2Δ-not4Δ-dhh1Δ-lsm1Δ* group and also to some extent in *pop2Δ* and *ccr4Δ* strains mRNAs normally transcriptionally activated during heat stress were slightly stabilized, while the transcriptionally silenced ones had modestly decreased abundance (Fig. 6b). Generally the mRNA level changes in each group were not greater than two-fold, explaining why most did not enter into the list of differentially expressed mRNAs. The mean pA-tail length of both gene classes was shorter in the core Ccr4-NOT mutants, particularly in the up-regulated group (Fig. 6b). Thus, the Not1-AID-*not2Δ-not4Δ-dhh1Δ-lsm1Δ* strain cluster seems to display some common stress-like response phenotype, though not acute enough to surpass a critical threshold.

Using RT-qPCR we have confirmed that the heat shock mRNAs *SSA4* and *HSP104* were up-regulated in the *not2Δ-not4Δ-dhh1Δ-lsm1Δ* strains, and that was associated with marked shortening of their pA-tails in the DRS datasets (from -11 to -21 adenosines on average; Fig. 6c). Heat-stress mRNAs are induced in response to sudden elevation of growth temperature, and in conditions of exposure to low levels of thiolutin (Adams and Gross, 1991; Czarnocka-Cieciura et al., 2024). In wild-type, *HSP104* and *SSA4* levels rose markedly in response to both stimuli. Despite initial up-regulation in the mutant strains, this dynamic was largely maintained, except for *lsm1Δ* (Fig. 6d; Supplementary Figure 6b), supporting the notion that perturbation of Ccr4-NOT- and decapping-dependent mRNA decay shifts cells toward a mild stress-like state, but the expression status is not acute enough to dampen any further stronger response. Importantly, though clearly present, this mild-stress response cannot fully explain all the abundance and pA-tail length changes observed in the tested strains. On the contrary, it opens the possibility that the heat shock mRNAs are in fact primary targets of the Not1-2-4-Lsm1-Dhh1 functional assembly.

## DISCUSSION

Accepted models for mRNA cytoplasmic turnover posit that considerable deadenylation of the 3’ polyadenosine tail is required for subsequent 5’end mRNA decapping and decay. Early works developed this model not surprisingly, as in general deadenylation is observed earlier during the mRNA cytoplasmic life, compared to decapping, leading to natural speculations about causality. Later works have pointed towards functional links between deadenylation and both the cytoplasmic polyA-binding protein Pab1/PABPC1 and the rate of mRNA translation, which were expected to collaborate to influence the onset of decapping (Caponigro and Parker, 1995; Funakoshi et al., 2007; Passmore and Coller, 2021). Physical links of the core Ccr4-NOT complex to decapping co-factors seemed to only reinforce the developing model (Maillet et al., 2001; Ozgur et al., 2010; Ukleja et al., 2016). However, several published reports downplay the role of deadenylation as a general trigger for mRNA degradation on reporter or larger groups of transcripts, showing that in the vast collection of cellular mRNAs, decay mechanisms of each have to be considered individually and in the context of response to environmental growth conditions (Muhlrad and Parker, 2005; Czarnocka-Cieciura et al., 2024; Audebert et al., 2024).

The cell possesses two main deadenylase complexes. The Pan2/3 complex binds directly to the pA-tail and to the polyA-binding protein; a binding mode that does not provide substrate specificity (Schäfer et al., 2014; Wolf et al., 2014). In contrast, the Ccr4-NOT is a much larger multi-subunit complex. Apart from direct association of its deadenylase subunits with RNA, the non-catalytic Not5 subunit also associates with the substrate in the context of translation, while the scaffolding Not1/CNOT1 protein provides extra support. In addition, the Ccr4-NOT interacts with a collection of six co-factors, the Puf1-6 proteins, which enhance recruitment to various substrates, and with the decapping co-factors (Wang et al., 2018). Therefore, it is not surprising that due to those properties, the Ccr4-NOT was found to target more efficiently mRNAs of low abundance, compared to Pan2/3 (Tudek et al., 2021). Taken together, it is most likely that the core non-catalytic subunits are the source of the Ccr4-NOT substrate specificity. Since the core displays links not only to the Puf1-6 proteins, but also to decapping factors, the natural question about the core’s true biological role arises.

Here, we have attempted to address the function of the Ccr4-NOT core using high-throughput transcriptomic approaches, DRS and Illumina, which were directed at describing the changes in mRNA abundance and polyadenylation profiles. The yeast model proved to be invaluable for this task, as we showed that, in contrast to the human system, it is dispensable for deadenylation. This functional shift was linked primarily to a change in the relative contribution to deadenylation of Ccr4 and Pop2 deadenylases in each organism. Pop2 provides a bridge between the Ccr4 and Not1 scaffold (Basquin et al., 2012). In human cells, it is the Pop2 orthologs that mediate most deadenylation (Yi et al., 2018), and unsurprisingly, the core CNOT1 is also required for deadenylation (Krawczyk et al., 2025). In contrast, most enzymatic activity in yeast is performed by Ccr4 (Tudek et al., 2021; Czarnocka-Cieciura et al., 2024); hence the role of Not1 is diminished. Ccr4 does not require the core NOT for deadenylation, as we have observed that its detachment from it, either by deletion of *POP2* or using a Not1 point mutant incapable of recruiting Pop2 (Basquin et al., 2012), does not elongate mRNA pA-tails the same way as *CCR4* deletion.

The possibility of detachment of the deadenylase function from the Ccr4-NOT core in yeast allowed phenotypic clustering of the core and catalytic subunits. This revealed global shortening of mRNA mean pA-tail length in core mutants, which was a phenotype previously specific to impairment in the decapping Dcp2 enzyme and in 5’-3’ exoribonuclease Xrn1 (Czarnocka-Cieciura et al., 2024). Accordingly, in this work we observed a similar phenotype in decapping co-factor mutants *lsm1Δ* and *dhh1Δ*. Supposedly, shorter mean pA-tails arise on mRNAs when their cytoplasmic dwelling time is extended (Audebert et al., 2024). That is why we also systematically evaluated changes to mRNA abundance. To this end, we first used the DRS dataset, which showed a clear correlation between pA-tail shortening and mRNA stabilization in *not2Δ*, *not4Δ* and Not1-AID strains, similar to *lsm1Δ* and *dhh1Δ*. In the case of *not3Δ* and *not5Δ* strains, a clear correlation was not observed, though many mRNAs did have a tendency to being up-regulated. The DRS dataset has its technical limitations, due to reduction in detection of mRNAs with very short pA-tails, which results from selection of the polyadenylated fraction prior to library synthesis (Tudek et al., 2021). Thus, considerable shortening of certain mRNA pA-tails could exclude their incorporation into the library, erroneously classifying them as down-regulated. If changes were global, this could also indirectly distort the abundance estimates of other mRNAs. Therefore, to more accurately determine changes to mRNA, we turned to Illumina sequencing. This again showed that the most important changes to mRNA abundance were characteristic of strains with impaired *NOT1*, *NOT2* and *NOT4* functions. These clustered closely with decapping mutants, *lsm1Δ* and *dhh1Δ*, but also with cells with depleted decapping enzyme Dcp2. Changes to mRNA abundance in other non-catalytic Ccr4-NOT subunits were somehow more limited compared to DRS, likely indicating an indirect impact of pA-tail shortening on abundance estimates.

In sum, this three-tier analysis allowed us to show that the Not1, Not2 and Not4 non-catalytic subunits of the Ccr4-NOT complex likely collaborate with the decapping factors Dhh1 and Lsm1 to regulate mRNA abundance. Importantly, we have ruled out the possibility that the phenotypic similarities are indirect and due to activation of a common gene expression pathway. The most consistent observed trend was associated with activation of the heat stress response, which is triggered by cytoplasmic protein misfolding. However, even though the heat shock protein encoding mRNAs were consistently up-regulated in *NOT1-2-4* and decapping co-factor mutants, resonating with previous reports for *not4Δ* (Preissler et al., 2015), the magnitude of the up-regulation did not match the one observed in *bona fide* stress conditions, nor did it preclude further up-regulation when a stress trigger was applied. It rather seems that *HSP* mRNAs are an important target of the Not1-2-4-Dhh1-Lsm1 functional assembly, and that their considerable pA-tail shortening and up-regulation in the mutants arise from delayed decapping.

Determining the role of the deadenylases, Ccr4 and Pop2, in regulation of decapping by the Not1-2-4 subunits is somewhat more challenging. At first, it is seemingly surprising that loss of *POP2* results in a more prominent growth defect compared to *CCR4* deletion. However, we observed that in the *pop2Δ* mutant, the overall changes to mRNA abundance are not only greater than those observed in *ccr4Δ*, but most strongly correlated with loss of *DHH1* or *LSM1*. Also, loss of the decapping co-factor proteins results in growth defects comparable to deletion of *POP2*. Thus, it is likely that Pop2 collaborates more tightly with the core NOT factors in regulating decapping than it does with Ccr4 in mediating deadenylation. Based on the direct interaction of Not1 with the Dhh1 decapping co-factor, previous studies proposed a model of mRNA handover from deadenylation to decapping (Maillet et al., 2001; Ozgur et al., 2010; Ukleja et al., 2016). However, in the high-throughput data we did not observe any sign supporting this model as a general rule. On the contrary, the mRNAs that were most susceptible to deadenylation by Ccr4-Pop2 were not the ones with the shortest pA-tails or increased abundance in *NOT1-2-4*-deficient strains, suggesting substrate separation. That said, the coupling might occur for certain targets, but is likely limited.

In sum, it seems reasonable to propose a more stochastic model explaining the relationship between deadenylation and decapping. Only a double *CCR4* and *PAN2* deletion of the two major deadenylases confers a strong proliferative defect in yeast and is associated with near-complete inhibition of deadenylation (Czarnocka-Cieciura et al., 2024; Tucker et al, 2001). Although single-deletion analyses indicate distinct substrate preferences, with Pan2/3 preferentially targeting highly abundant mRNAs and Ccr4 being more active on less abundant transcripts, the severe phenotype of the double mutant demonstrates substantial functional redundancy between the two deadenylase pathways (Tudek et al., 2021; Czarnocka-Cieciura et al., 2024). Thus, in the absence of one deadenylase, the other can partially compensate, ultimately sustaining viability. The trigger for mRNA decapping may therefore be multifactorial, reflecting the integration of several features of the mRNP, including its translational status and ribosome occupancy, its overall structural organization, and the occupancy of the polyA-tail binding proteins (Caponigro and Parker, 1995; Tsuboi and Inada, 2010; Huch and Nissan, 2014; Franks and Lykke-Andersen, 2008). Thus, mRNAs might enter the pool of eligible substrates for decapping only after minimal random deadenylation occurs, and at this point are subject to stochastic probing by decapping co-factors, which each are sensitive to particular features of mRNA (He and Jacobson, 2023; Franks and Lykke-Andersen, 2008; Labeauvie et al., 2026). Our data suggest that the core Ccr4-NOT complex, at least in yeast, is one among many such decapping co-factors. The role of Pop2 is dual; it can provide limited deadenylation activity, but it also works closely with the core, allowing it to express its role in decapping.

## METHODS

### Yeast and HEK293 cell line growth conditions

Yeast cultures were grown in YPDA media at temperatures indicated for each experiment. Treatment with thiolutin (Sigma; T3450) was performed by adding the compound to a final concentration of 4 ug/ml. Samples for all chase experiments were collected by mixing the cell culture with an equal volume of ice-cold ethanol, which inactivates cellular metabolism. To deplete Not1 using an AID tag (Nishimura et al, 2009; Morawska and Ulrich, 2013), 1-Naphthaleneacetic acid (1-NAA; N0640-25G, Merck) or auxin sodium salt (IAA; Indole-3-acetic acid sodium salt; I5148-2G; Merck) was added to a final concentration of 1 mM. The heat stress chase experiment was performed by pre-culturing cells at 25 °C and adding an equal volume of media pre-heated to 51 °C, resulting in a final temperature of 37 °C, which was maintained by culture incubation in a water bath. The list of yeast strains used in this study is given in Supplementary Table 1.

HEK293 Flp-In T-REx cells (R78007; Thermo Fisher Scientific) were grown in Dulbecco’s Modified Eagle Medium (DMEM) with high glucose (392-0415; VWR) supplemented with 10% Fetal Bovine Serum (FBS) (A5256701 or 10270106; Life Technologies) at 37 °C with 5% CO_2_. To activate expression of microRNAs directed at CNOT1/6/6L/7/8, cells were treated with tetracycline to a final concentration of 100 ng/ml for three days.

### Preparation of stable HEK293 cell lines expressing microRNAs directed against CNOT-encoding mRNAs

The microRNAs directed against mRNAs encoding CNOT1, CNOT6, CNOT6L, CNOT7 and CNOT8 were designed using the BLOCK-iT™ RNAi Designer (https://rnaidesigner.thermofisher.com/rnaiexpress/). The selection criteria included a GC content of 35–55% and mapping of the miRNAs to an open reading frame. To enhance knockdown efficiency, one construct contained three miRNAs targeting different regions of the mRNA sequence. The sequences of the miRNA cassettes are provided in Supplementary Table 2. The constructs were cloned into the acceptor vector containing a bi-directional promoter activated by tetracycline, pKK-B16 *BspT*I_*Not*I_mCherry/EGFP_*BshT*I_*Nhe*I. The obtained plasmids are listed in Supplementary Table 3.

### Yeast and human proliferation assays

To evaluate yeast growth, cells were spotted on YPDA plates in serial 10x dilutions starting from OD ∼0.4 and grown at designated temperatures for 2-3 days.

To assess HEK293 cell line growth rates, we used the parental HEK293 Flp-In TREx cell line and its derivatives containing genome-integrated plasmids encoding microRNAs targeting CNOT mRNAs. MicroRNA expression was induced by treatment with tetracycline at a final concentration of 100 ng/mL for a total of 6 days. Briefly, cells were initially cultured on 60-mm dishes in the presence of tetracycline for 3 days. They were then transferred to 96-well plates and cultured for an additional 3 days in an IncuCyte S3 live-cell imaging system (4647; Essen BioScience), where cell growth was monitored. Roughly 5000 cells (targeting initial confluence below 20 %) were seeded per well on a 96-well plate. At least 4 to 5 wells were prepared as technical replicates to estimate confluence changes over time. Growth was evaluated in 4-6 hour intervals by taking one bright-field picture of each well. Confluence data was analyzed using the IncuCyte software (version 2019B Rev2).

To quantitatively analyze the HEK293 cell line growth, logistic fitting to raw confluence data was performed using the nls protocol in RStudio (version 2024.04.2+764), as previously described in Tomecki et al. (2026).

### RNA extraction

Total RNA was isolated using the hot acid phenol extraction method. Cell pellets were resuspended in 400 μl of phenol solution equilibrated with 0.1 M citrate buffer at pH 4.3 (Sigma; P4682), followed by the addition of 400 μl TES buffer (10 mM Tris pH 7.5, 5 mM EDTA, 1% SDS). The samples were vortexed for 45 min at 65 °C and subsequently centrifuged at 4 °C for 10 min. The resulting supernatant was transferred to a new tube, and an additional 400 μl of phenol solution equilibrated with 0.1 M citrate at pH 4.3 was added. Samples were vortexed again for 20 min at 65 °C and centrifuged at 4 °C for 10 min. The aqueous phase was then transferred to a fresh tube and mixed with 400 μl chloroform (C2432; Sigma). Samples were briefly vortexed at room temperature to facilitate phenol removal and centrifuged at 4 °C for 10 min.

Following centrifugation, the supernatant was transferred to a clean tube, and 45 μl of 2 M LiCl together with 1 ml of 95% ethanol was added. RNA was precipitated at −80 °C for a minimum of 30 min and then centrifuged at 4 °C for 25 min. The resulting pellets were washed with 400 μl of 80% ethanol and, after removal of the supernatant, dried at 37 °C. RNA pellets were subsequently dissolved in nuclease-free water, and RNA concentration was determined using a NanoDrop spectrophotometer.

### Reverse-transcription and quantitative PCR

To perform reverse transcription, the RNA samples were first treated with DNase using the TURBO DNA-free™ Kit Quantity (AM1907; Life Technologies). 5 μg of RNA was used in a total reaction volume of 12 μl according to the manufacturer’s instructions. 1-2 μg of DNase-treated RNA was used to perform reverse transcription using the SuperScript IV (18090050; Life Technologies) and a mix of oligo dT and random hexamers (N8080127; Life Technologies) following the manufacturer’s instructions. The reaction was diluted to 400 μl and used as template for qPCR. In case of primers specific to yeast 25S rRNA, the template was further diluted 1:200. Quantitative PCR reactions were prepared using the RT PCR Mix SYBR® kit (2008-1000; A&A Biotechnology), and reactions were run on a Roche LightCycler 480 machine. Data were collected using the 2^nd^ derivative max protocol. The list of oligonucleotides used in this study is given in Supplementary Table 4.

### Purification of polyadenylated RNA for preparation of DRS libraries

The poly(A)+ RNA fraction was isolated using magnetic beads conjugated to oligo-dT (Life Technologies; 61005). Briefly, 35 μg of total RNA was dissolved in 50 μl of nuclease-free water and combined with an equal volume of binding buffer (20 mM Tris-HCl, pH 7.5, 1 M LiCl, 2 mM EDTA). The RNA was denatured at 65 °C for 2 min and subsequently placed on ice. For each 35 μg of total RNA, 100 μl of bead slurry was used. The beads were pre-washed three times with 1 ml of binding buffer and then resuspended in 50 μl of binding buffer per sample.

The washed beads were added to the denatured RNA and incubated at room temperature for 20 min with intermittent mixing. Following incubation, the supernatant was removed, and the beads were washed twice with 100 μl of wash buffer (10 mM Tris-HCl, pH 7.5, 150 mM LiCl, 1 mM EDTA). After removing residual wash buffer, the beads were resuspended in 12 μl of nuclease-free water. The samples were then incubated at 80 °C for 2 min, after which the supernatant containing the eluted poly(A)+ RNA was collected and used for sequencing library preparation.

### Bulk pA-tail length analysis

For 3′-end labeling, 500 ng of oligo(dT)-selected poly(A)+ RNA was incubated in a 20 μl reaction mixture containing 2 μl of 10× Reaction Buffer for T4 RNA Ligase (Thermo Scientific), 2 μl of 10 mM ATP, 2 μl of 100 mM DTT (Invitrogen), 2 μl of 100% DMSO (Thermo Scientific), 0.5 μl of RiboLock RNase Inhibitor (40 U/μl; Thermo Scientific), 2 μl of T4 RNA Ligase 1 (EL0021; Thermo Scientific), and 1.5 μl of [5′-32P]pCp (3000 Ci/mmol, 10 mCi/ml; Hartmann Analytic). The reaction was carried out for 1 h at 37 °C, after which the enzyme was heat-inactivated at 95 °C for 5 min. The labeled RNA was subsequently digested with 0.5 μg of RNase A and 10 U of RNase T1 (both Thermo Scientific) for 20 min at 37 °C. The reaction was terminated by adding an equal volume of formamide loading dye (90 % formamide in 1× TBE, 0.03 % xylene cyanol, 0.03 % bromophenol blue), followed by incubation at 95 °C for 5 min and immediate freezing in liquid nitrogen. A 5 μl aliquot of each sample was separated on a denaturing 12% polyacrylamide sequencing gel containing 8 M urea in 1× TBE. As molecular size markers, 3′-end-labeled synthetic RNA oligonucleotides of 20, 31, and 51 nt in length (CCCCACCACCAUCACUUA3, CCCCACCACCAUCACUUA14, and CCCCACCACCAUCACUUA34, respectively; Future Synthesis) were electrophoresed alongside the labeled poly(A) tails. After electrophoresis, the gel was transferred onto Whatman 3MM filter paper and dried under vacuum at 80 °C for 2 h. The dried gel was then exposed overnight to a PhosphorImager screen (FujiFilm) and scanned using an FLA 9000 scanner. The autoradiogram was quantified using ImageJ (Fiji; version 2.14/154f; build c89e8500e4), and visualized using the ggplot2 package from RStudio (version 2024.04.2+764).

### Nanopore sequencing

DRS libraries were prepared using the Direct RNA Sequencing Kit (ONT—Oxford Nanopore Technologies, SQK-RNA002) following the manufacturer’s protocol. Briefly, 50–200 ng of oligo-dT(25)-enriched mRNA isolated from *Saccharomyces cerevisiae* yeast was combined with cap-enriched or total RNA derived from other organisms, including human, mouse, *A. thaliana*, and *C. elegans*, as previously described by Bilska et al. (2020). Raw sequencing data were base called using Guppy (ONT).

### Whole cell extract preparation and western blot analysis

For preparation of yeast whole cell extracts, cells corresponding to 4–6 OD units were resuspended in 8 M urea and incubated at 80 °C for 5 minutes. Glass beads were then added, and cells were disrupted by vortexing for 5 min. The urea extracts were clarified by centrifugation, and the supernatants were collected. Protein concentrations were determined, and extracts were adjusted to equal protein concentrations before resuspension in Laemmli buffer. A total of 30 μg of protein was loaded onto 6 % or 10 % denaturing SDS-PAGE gels.

For preparation of HEK293 cell line extracts cell pellets were resuspended in 40–80 μl of RIPA buffer (50 mM Tris-HCl, pH 8.0, 150 mM NaCl, 1 % NP-40, 0.5 % sodium deoxycholate, and 0.1 % SDS) supplemented with 1 mM PMSF and a protease inhibitor cocktail (2 μM pepstatin A, 0.6 μM leupeptin, 2 mM benzamidine, and 2 μg/ml chymostatin). Samples were sonicated using a Bioruptor XL sonicator (Diagenode) at high power for 20 cycles of 15 s on and 15 s off. Lysates were then centrifuged at 20,000 × g for 15 min at 4 °C, and the supernatants were collected. Supernatants were mixed with Laemmli buffer and analyzed by SDS-PAGE as described above.

Following electrophoresis, proteins were transferred to Amersham nitrocellulose Protran Western blotting membranes (GE10600001) using a semi-dry transfer system (Bio-Rad, cat. no. 1703940). Membranes were blocked in TBS-T containing 5 % milk and incubated overnight with the appropriate primary antibodies. The antibodies used were anti-FLAG (F1804, Sigma), anti-Pgk1 (Novex Life Technologies; discontinued), anti-CNOT1 (D5M1K; 44613; Cell Signaling), anti-CNOT7 (D3M6O; 86665; Cell Signalling), and anti-GAPDH (GA1R; MA515738; LifeTechnologies). Following washing with TBS-T, membranes were incubated for 1 h with the appropriate HRP-conjugated secondary antibodies in TBS-T containing 5 % milk. Membranes were subsequently washed and developed using Clarity Western ECL substrate (Bio-Rad) using Fujifilm RX-N x-ray films.

### Illumina library construction and processing of raw data

3′ mRNA-seq libraries were prepared using an anchored oligo dT provided by the QuantSeq 3′ mRNA-Seq V2 Library Prep Kit FWD for Illumina (Lexogen, cat. no. 015.96) and Lexogen UDI 12 nt Unique Dual Indexing Set A1 (cat. no. 101.96), following the manufacturer’s instructions. Libraries were amplified by 13 PCR cycles, assessed for fragment size using an Agilent TapeStation 4150 with High Sensitivity D1000 reagents, and quantified by qPCR using the KAPA Library Quantification Kit. Sequencing was performed on an Illumina NovaSeq 6000 using the S1 Reagent Kit v1.5 (200 cycles), with paired-end reads (2 × 100 cycles) and a 1 % PhiX control spike-in.

Raw sequences were trimmed according to quality using Trimmomatic (Bolger, Lohse, and Usadel, 2014) (version 0.39) using default parameters, except: SLIDINGWINDOW:4:20 MINLEN:20. Trimmed sequences were mapped to the *S. cerevisiae* reference genome previously reported in Tudek et al. (2021) (version S288C) using Hisat2 (Kim, Langmead, and Salzberg, 2015) with default parameters. Optical duplicates were removed using the MarkDuplicates tool from the GATK (McKenna et al., 2010) package (version 4.2.3.0) with default parameters except OPTICAL_DUPLICATE_PIXEL_DISTANCE set to 12000. Reads that failed to map to the reference were extracted using Samtools (Li et al., 2009) and mapped to the Silva meta-database of rRNA sequences (Quast et al., 2013) (version 119) with Sortmerna (Kopylova, Noé, and Touzet, 2012) (version 2.1b) using the “–best 1” option. Mapped reads were associated with transcripts from the User-provided database using HTSeq-count (Anders, Pyl, and Huber, 2015) (version 2.0.1) with default parameters. Differentially expressed genes were selected using the DESeq2 package (Love, Huber, and Anders, 2014) (version 1.16.1). Fold change was corrected using apeglm (Zhu, Ibrahim, and Love, 2019).

### Bioinformatic analysis of DRS and Illumina data

Statistics regarding DRS and Illumina datasets, such as fold-change in mRNA level, mean and median pA-tail lengths, were summarized in tables deposited in the Mendeley Data repository (doi 10.17632/bp6yysczxk.1), along with the R code underlying each figure. Primary statistics are listed in Supplementary Tables 5 and 6. The data were further analyzed and visualized using RStudio (version 2024.04.2+764) using custom scripts and the ggplot2 and corrplot packages. For each strain or depletion condition, transcript-level changes in pA-tail length were calculated relative to the corresponding wild-type or control condition. While the Illumina dataset presents one collection of detected RNAs, each DRS dataset is given separately; that is why, to cross-compare the DRS datasets, transcript identifiers were merged across datasets to generate a common transcript-level matrix. The merged dataset was deduplicated, and a single entry corresponding to *YGL076C* mRNA was removed as it was misannotated in the original reference. As a rule, only mRNAs represented by at least 10 reads for each DRS dataset were analyzed.

The distribution of transcript-level pA-tail length changes was visualized using a collection of density plots using the ggplot2 geom_density function, with zero representing no change relative to the corresponding control.

To produce correlation matrices of pA-tail length or abundance change, first a matrix of Spearman ρ correlation coefficients was prepared using the pairwise complete observation setting so that transcripts lacking measurements in one condition could still contribute to comparisons for which measurements were available. The resulting correlation matrix was visualized using hierarchical clustering and correlation plots. Hierarchical clustering was performed using complete linkage on a distance matrix defined as 1-ρ. Pairwise sample sizes were also calculated for every comparison to document the number of transcripts contributing to each correlation estimate.

For selected pairs of conditions, scatter plots with two-dimensional density contours and linear regression lines were generated using the ggplot2 package (using the geom_point, geom_density_2d_filled, and geom_smooth “lm” functions).

To identify pronounced pA-tail length changes, a 5-adenosine mean difference with respect to the wild-type reference was considered. To identify mRNAs with significantly changed abundance, a two-fold difference in levels compared to the wild-type reference was selected. To identify the transcripts with changed features in each set, the intersect function in RStudio was used, selecting the “transcript” column. To test whether the observed overlaps between transcript groups were greater or smaller than expected from the available transcript universe, Fisher’s exact tests were performed using 2 × 2 contingency tables. The universe was restricted to the transcripts for which measurements were available for all conditions included in these analyses. Odds ratios and exact p-values were calculated using Fisher’s exact test with the alternative hypothesis specified as a negative association. This analysis was used to assess the statistical significance of the overlap between transcript groups rather than relying solely on the observed intersection size.

For visualization of transcript abundance changes, the heatmap function in RStudio was used. Extreme values were limited to a range of −3 to +3 or -2.5 to +2.5, depending on the RNA type. This was done to prevent outliers from dominating the colour scale. Heatmaps were generated using a blue–white–red colour scale, with blue and red representing decreased and increased transcript abundance, respectively. Perturbations were hierarchically clustered according to the similarity of their transcriptome-wide responses. Pairwise Spearman correlations were calculated between perturbations, and correlation distances were defined as 1 − ◻. Hierarchical clustering was performed using average linkage, and the resulting dendrogram was used to order the columns of the heatmaps. In a complementary analysis, transcripts were ordered according to normalized WT expression levels rather than clustered, allowing abundance-dependent response patterns to be visualized. The same correlation-based hierarchical clustering approach was applied to pA-tail-length changes, using pairwise Spearman correlations calculated from transcripts shared between datasets.

## DATA AVAILABILITY

Raw data underlying this study are available from the corresponding authors upon reasonable request.

## AUTHOR CONTRIBUTIONS

AT conceptualized the project, prepared most yeast cultures for sequencing, prepared yeast growth assays, validated depletion of Not1-AID by western blot, validated CNOT mRNA down-regulation by RT-qPCR, performed numerical analysis of HEK293 growth, analyzed mRNA expression changes in yeast by RT-qPCR, performed bioinformatical analysis of the processed DRS and Illumina data, wrote the first draft, prepared figures and raw data deposition, obtained funding. RT curated sample preparation for Illumina sequencing, obtained stable HEK293 cell lines, validated depletion of CNOT1 and CNOT7 by western blot, evaluated bulk pA-tail length distributions in CNOT-depleted cells. AJ prepared the HEK293 proliferation assays and edited the manuscript. MT obtained the Not1-AID strain and edited the manuscript. AP performed the *not2Δ*, *not4Δ*, *lsm1Δ* and *dhh1Δ* heat stress analysis. KG analyzed raw Illumina sequencing data. PK and ACC analyzed raw DRS sequencing data. SM supervised DRS sequencing. AD obtained funding, supervised and conceptualized the project.

## ACKNOWLEDGMENTS / FUNDING INFORMATION

This work was supported by National Science Centre SONATA grant awarded to AT (2020/39/D/NZ2/02174).

## SUPPLEMENTARY INFORMATION

**Supplementary Figure 1.**
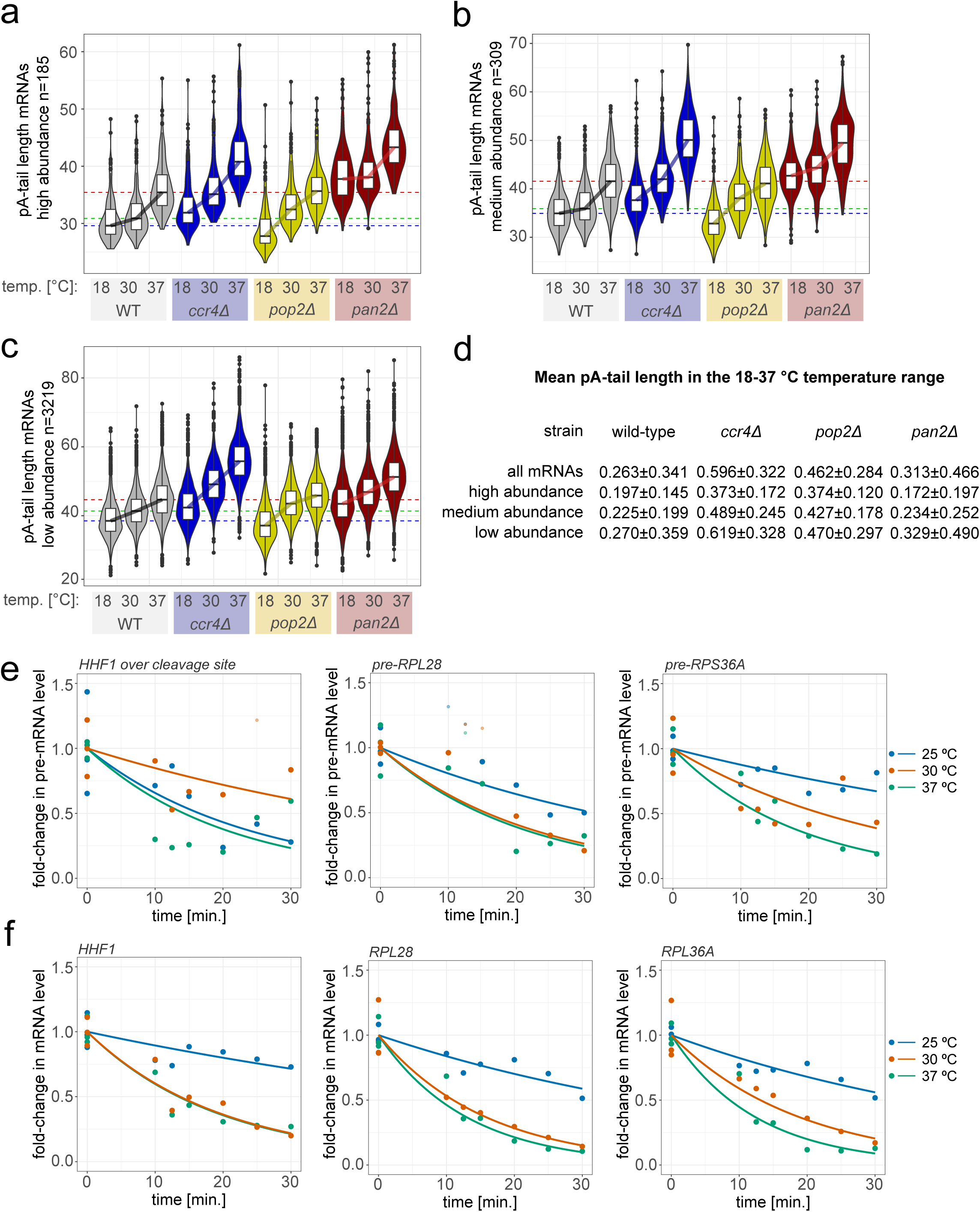
Ccr4-NOT dampens the temperature-dependent increase in pA-tail lengths. **a-c.** Box and violin plots illustrate the changes to mean mRNA pA-tail lengths for wild-type, *ccr4Δ*, *pop2Δ* and *pan2Δ* strains grown continuously at 18, 30 and 37 °C for mRNAs of high (a), medium (b) and low abundance (c). **d.** The table presents the numerical data underlying the graph presented in Fig. 1b, that is, the mean increase and standard deviation in individual mRNA pA-tail length calculated for cells grown at 18/30 and 18/37 °C. **e-f.** Time-course RT-qPCR analysis of the levels or the *RPL28*, *RPL36A*, and *HHF1* mRNA precursors using primers spanning the exon and intron junction, for ribosome subunit mRNAs, or over the *HHF1* 3’end cleavage site (e), and the corresponding mature forms (f) in wild-type cells treated with thiolutin at 18, 30 or 37 °C. The single dots represent experimental measurements, while the trend line was inserted using the non-linear fitting protocol in R.

**Supplementary Figure 2.**
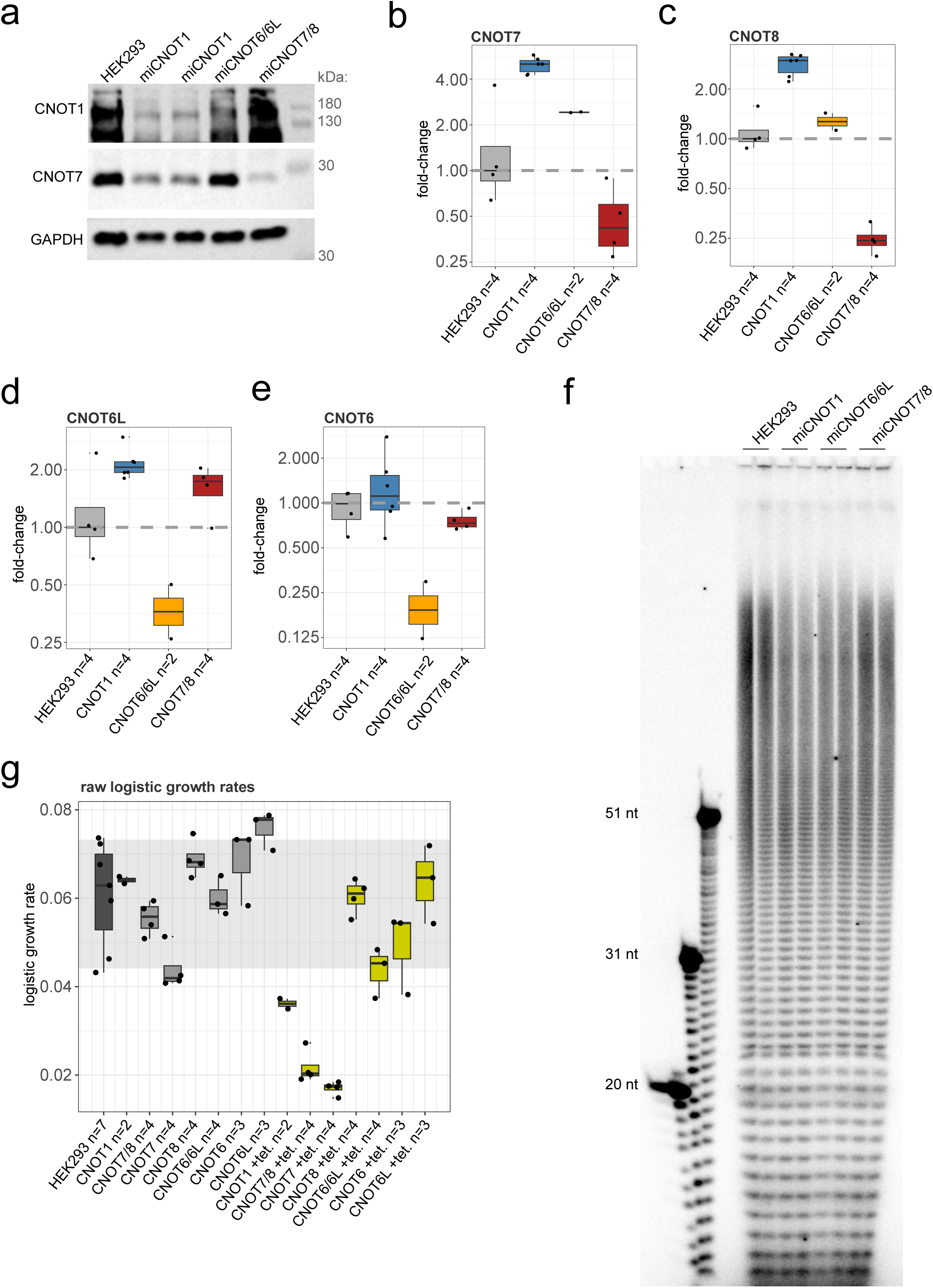
Particular Ccr4-NOT subunits retain their essential character, but diverge in deadenylation roles in yeast compared to human. **a.** Western blot analysis of knock-down efficiency of CNOT1 and CNOT7 in HEK293 control and cells expressing microRNAs against tested proteins. GAPDH was revealed as a loading control. **b-e.** RT-qPCR analysis of the levels of mRNAs coding for *CNOT6*, *CNOT6L*, *CNOT7*, and *CNOT8* in control HEK293 cells and those knocked down for CNOT1, CNOT6/6L and CNOT7/8. **f.** Image of an autoradiogram showing the analysis of pA-tail lengths in HEK293 control and cells knocked down for CNOT1, CNOT6/6L or CNOT7/8. Separately are loaded three molecular weight markers. The image quantification is shown in Fig. 2e. **g.** The box plot shows raw logistic growth rates calculated for control HEK293 and cells knocked down for CNOT1, CNOT6, CNOT6L, CNOT7, CNOT8, CNOT6/6L or CNOT7/8. The fold-change analysis between cells expressing microRNAs and not induced is shown in Fig. 2f.

**Supplementary Figure 3.**
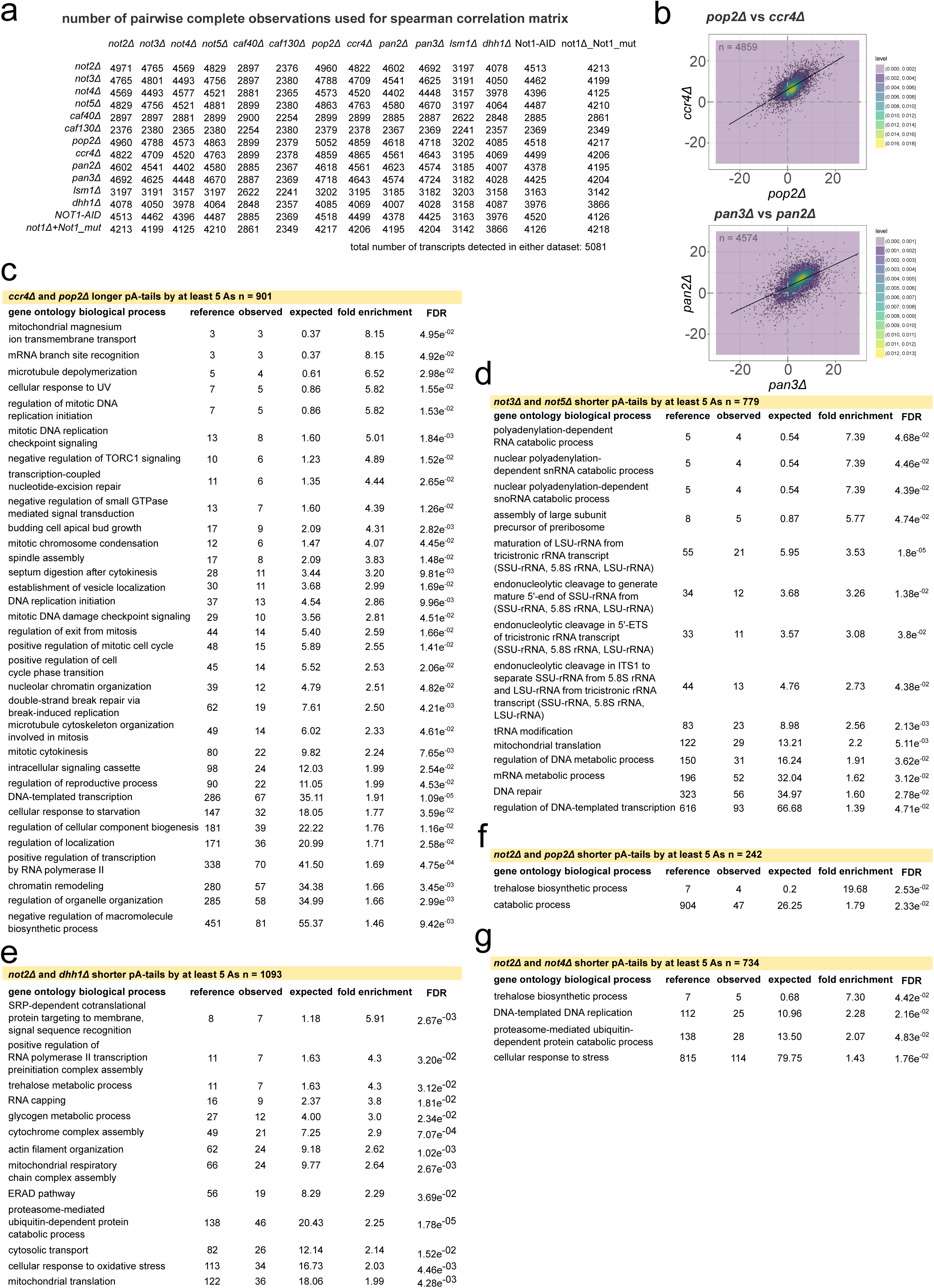
Loss of yeast core Ccr4-NOT subunits results in mRNA pA-tail length shortening akin to decapping mutants. **a.** Table shows the number of pairwise observations used to calculate Spearman ρ correlation coefficients shown in Fig. 3c. **b.** Scatter plots compare mRNA pA-tail length changes relative to wild-type between *pop2Δ*-*ccr4Δ* and *pan2Δ*-*pan3Δ*. The number of pairwise observations is indicated on each graph. A two-dimensional density plot is overlaid on top of the scatter, and a linear regression line is drawn through the data. The Spearman ρ correlation coefficients are shown in Fig. 3c. **c-g.** Tables list the main gene ontology groups for groups of mRNAs with elongated pA-tails in *pop2Δ*-*ccr4Δ* (c) or shortened pA-tails in *not3Δ-not5Δ (d), not2Δ*-*dhh1Δ (e), not2Δ*-*pop2Δ (f) and not2Δ*-*not4Δ* (g). Gene Ontology groups were constructed using the GeneOntology suite (https://geneontology.org/), choosing the biological process function.

**Supplementary Figure 4.**
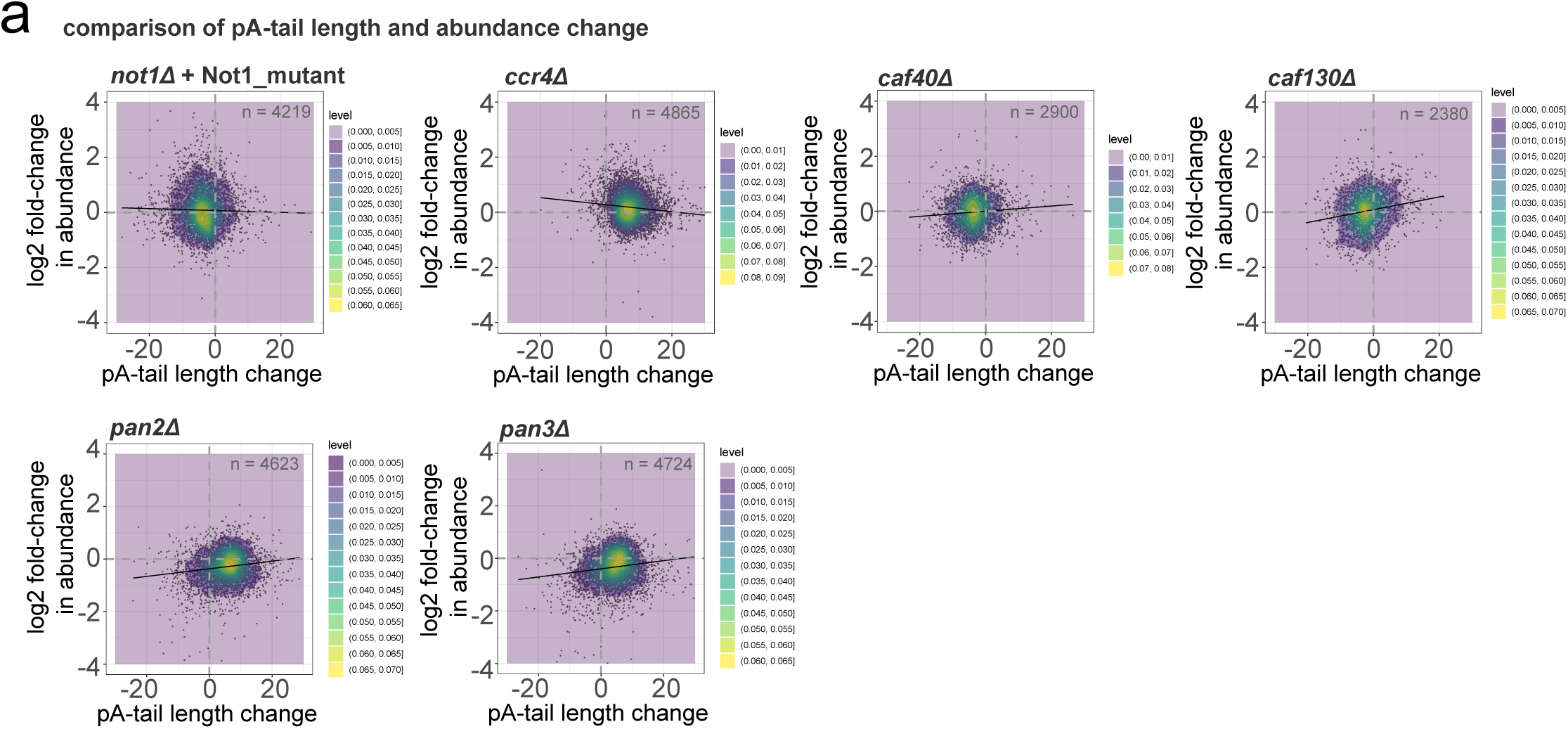
Loss of Not1-2-4 and decapping co-factors results in simultaneous pA-tail shortening and mRNA up-regulation. **a.** Scatter plots show pA-tail length and log2 abundance changes in mutant strains (*not1Δ_Not1_mut*, *cccr4Δ*, *caf40Δ*, *caf130Δ*, *pan2Δ*, *pan3Δ*) or in cells depleted of Not1 using AID system compared to wild-type. The number of mRNAs examined is given as ‘n =’. A linear regression line is drawn; the Spearman ρ coefficients are listed in Fig. 4b.

**Supplementary Figure 5.**
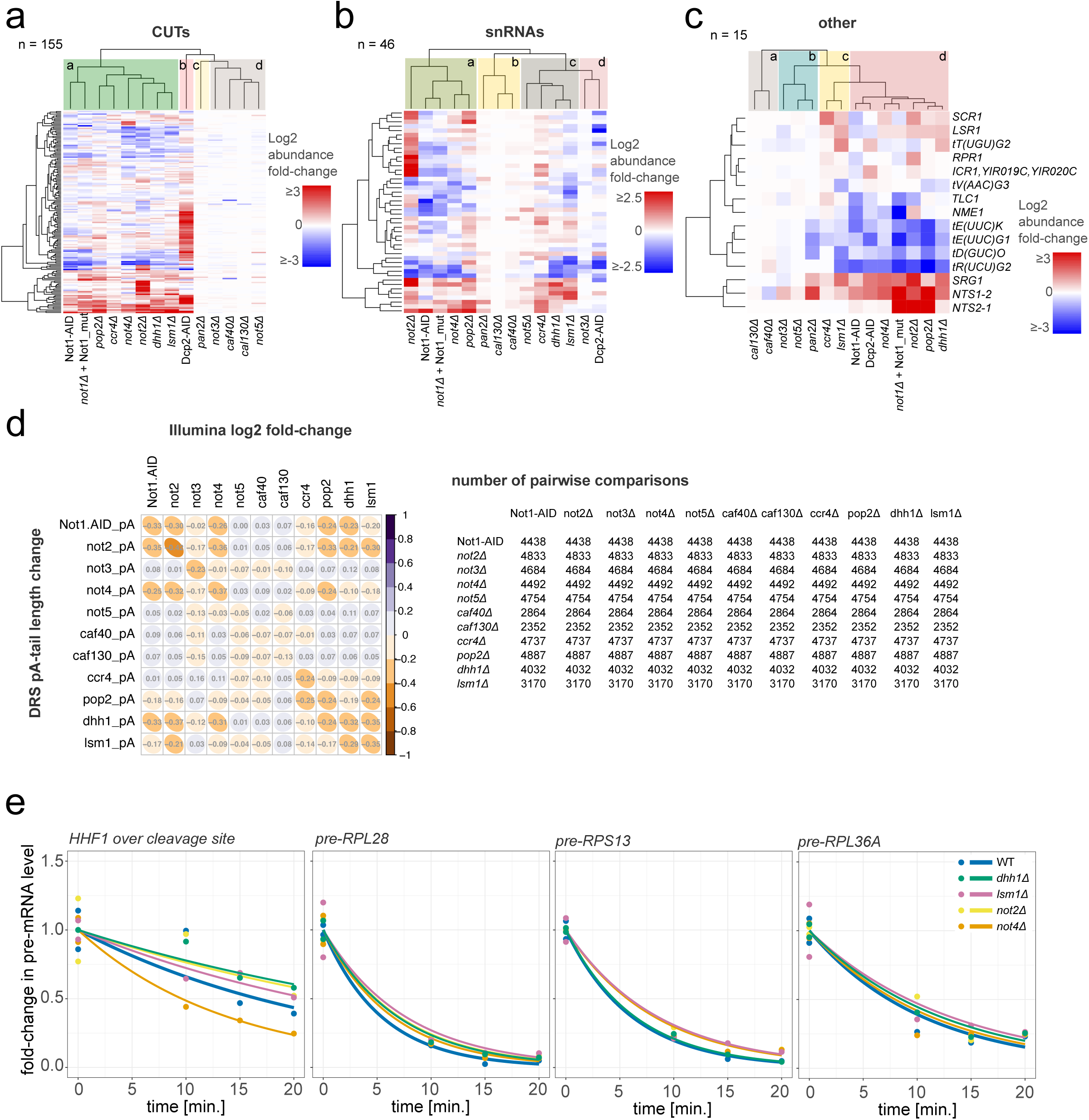
Loss of the Not1-2-4 core Ccr4-NOT subunits and Pop2 manifests phenotypic similarities to deletions of decapping co-factors *DHH1* and *LSM1*. **a-c.** Heat maps show CUTs (a), snRNAs (b), and other ncRNAs (c) abundance changes in oligo-dT-primed Illumina datasets derived from strains impaired for Dcp2, Dhh1, Lsm1, and Ccr4-NOT or Pan2/3 subunits relative to wild-type. The heat map is sorted by ncRNA expression similarity, while the strains are grouped by similarity of Spearman ρ coefficients. **d.** Matrix comparing Spearman ρ coefficients between mRNA pA-tail length in DRS and log2 abundance fold change in Illumina datasets. The table on the right shows the number of pairwise comparisons available to derive the coefficients. **e.** RT-qPCR analysis of nascent *HHF1*, *RPL28*, *RPS13*, and *RPL36A* levels in thiolutin time-course for wild-type, *not2Δ*, *not4Δ*, *dhh1Δ* and *lsm1Δ* strains. The abundance of the mature mRNA forms is shown in Fig. 5f.

**Supplementary Figure 6.**
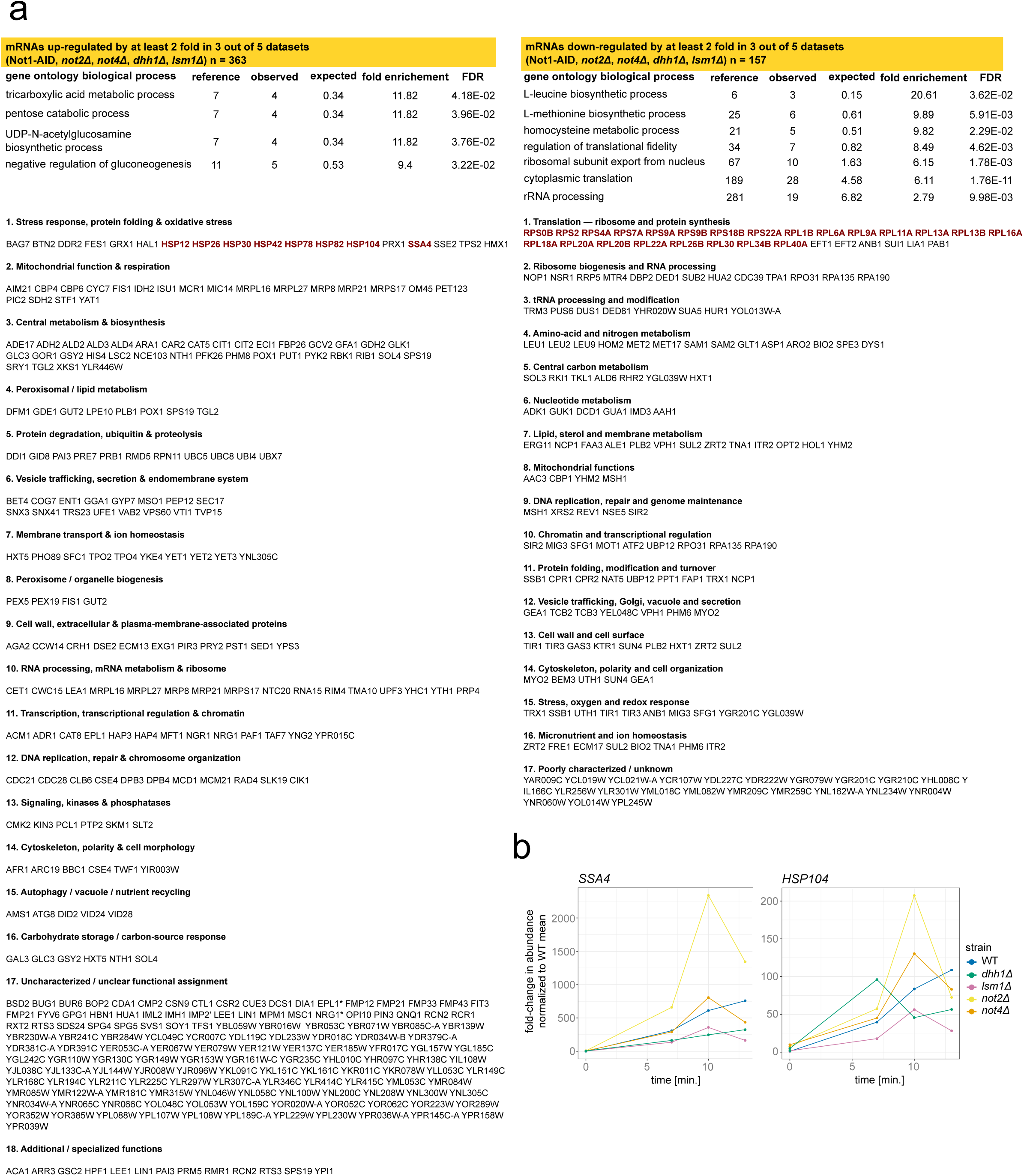
Loss of Not1-2-4 and Dhh1, Lsm1 induces a mild stress response. **a.** Top tables list the gene ontology groups detected for mRNAs up- or down-regulated in Not1-AID, *not2Δ*, *not4Δ*, *dhh1Δ* or *lsm1Δ*. Transcripts were selected if they were differentially expressed in 3 out of 5 examined datasets. The table at the bottom lists the affected mRNAs, grouping them functionally. In dark red are highlighted mRNAs transcriptionally impacted during the acute response to heat stress as defined by Vinayachandran et al. (2018). **b.** Graphs show *HSP104* and *SSA4* levels in wild-type, *not2Δ*, *not4Δ*, *dhh1Δ* or *lsm1Δ* strains, at 25 °C (time 0), and during three time-points sampled after sudden shift to growth at 37 °C measured by RT-qPCR.

**Supplementary Table 1.**
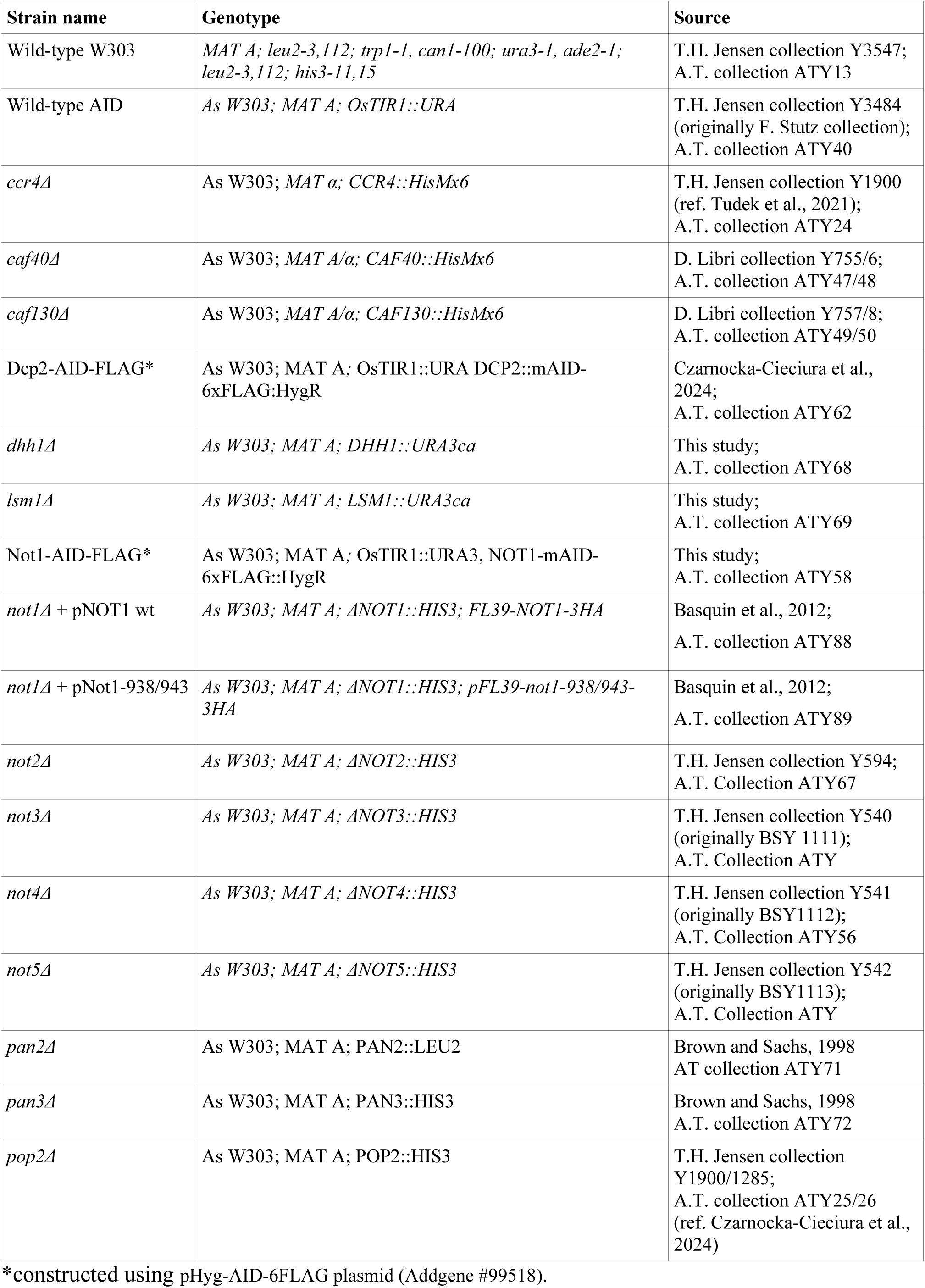
List of yeast strains used in this work.

| **Strain name** | **Genotype** | **Source** |
|---|---|---|
| Wild-type W303 | *MAT A; leu2-3,112; trp1-1, can1-100; ura3-1, ade2-1; leu2-3,112; his3-11,15* | T.H. Jensen collection Y3547;<br>A.T. collection ATY13 |
| Wild-type AID | *As W303; MAT A; OsTIR1::URA* | T.H. Jensen collection Y3484 (originally F. Stutz collection);<br>A.T. collection ATY40 |
| *ccr4Δ* | As W303; *MAT α; CCR4::HisMx6* | T.H. Jensen collection Y1900 (ref. Tudek et al., 2021);<br>A.T. collection ATY24 |
| *caf40Δ* | As W303; *MAT A/α; CAF40::HisMx6* | D. Libri collection Y755/6;<br>A.T. collection ATY47/48 |
| *caf130Δ* | As W303; *MAT A/α; CAF130::HisMx6* | D. Libri collection Y757/8;<br>A.T. collection ATY49/50 |
| Dcp2-AID-FLAG* | As W303; MAT A; OsTIR1::URA DCP2::mAID-6xFLAG:HygR | Czarnocka-Cieciura et al., 2024;<br>A.T. collection ATY62 |
| *dhh1Δ* | *As W303; MAT A; DHH1::URA3ca* | This study;<br>A.T. collection ATY68 |
| *lsm1Δ* | *As W303; MAT A; LSM1::URA3ca* | This study;<br>A.T. collection ATY69 |
| Not1-AID-FLAG* | As W303; MAT A; OsTIR1::URA3, NOT1-mAID-6xFLAG::HygR | This study;<br>A.T. collection ATY58 |
| *not1Δ* + pNOT1 wt | *As W303; MAT A; ΔNOT1::HIS3; FL39-NOT1-3HA* | Basquin et al., 2012;<br>A.T. collection ATY88 |
| *not1Δ* + pNot1-938/943 | *As W303; MAT A; ΔNOT1::HIS3; pFL39-not1-938/943-3HA* | Basquin et al., 2012;<br>A.T. collection ATY89 |
| *not2Δ* | *As W303; MAT A; ΔNOT2::HIS3* | T.H. Jensen collection Y594;<br>A.T. Collection ATY67 |
| *not3Δ* | *As W303; MAT A; ΔNOT3::HIS3* | T.H. Jensen collection Y540 (originally BSY 1111);<br>A.T. Collection ATY |
| *not4Δ* | *As W303; MAT A; ΔNOT4::HIS3* | T.H. Jensen collection Y541 (originally BSY1112);<br>A.T. Collection ATY56 |
| *not5Δ* | *As W303; MAT A; ΔNOT5::HIS3* | T.H. Jensen collection Y542 (originally BSY1113);<br>A.T. Collection ATY |
| *pan2Δ* | As W303; MAT A; PAN2::LEU2 | Brown and Sachs, 1998<br>AT collection ATY71 |
| *pan3Δ* | As W303; MAT A; PAN3::HIS3 | Brown and Sachs, 1998<br>A.T. collection ATY72 |
| *pop2Δ* | As W303; MAT A; POP2::HIS3 | T.H. Jensen collection Y1900/1285;<br>A.T. collection ATY25/26 (ref. Czarnocka-Cieciura et al., 2024) |

\*constructed using pHyg-AID-6FLAG plasmid (Addgene #99518).

**Supplementary Table 2.**
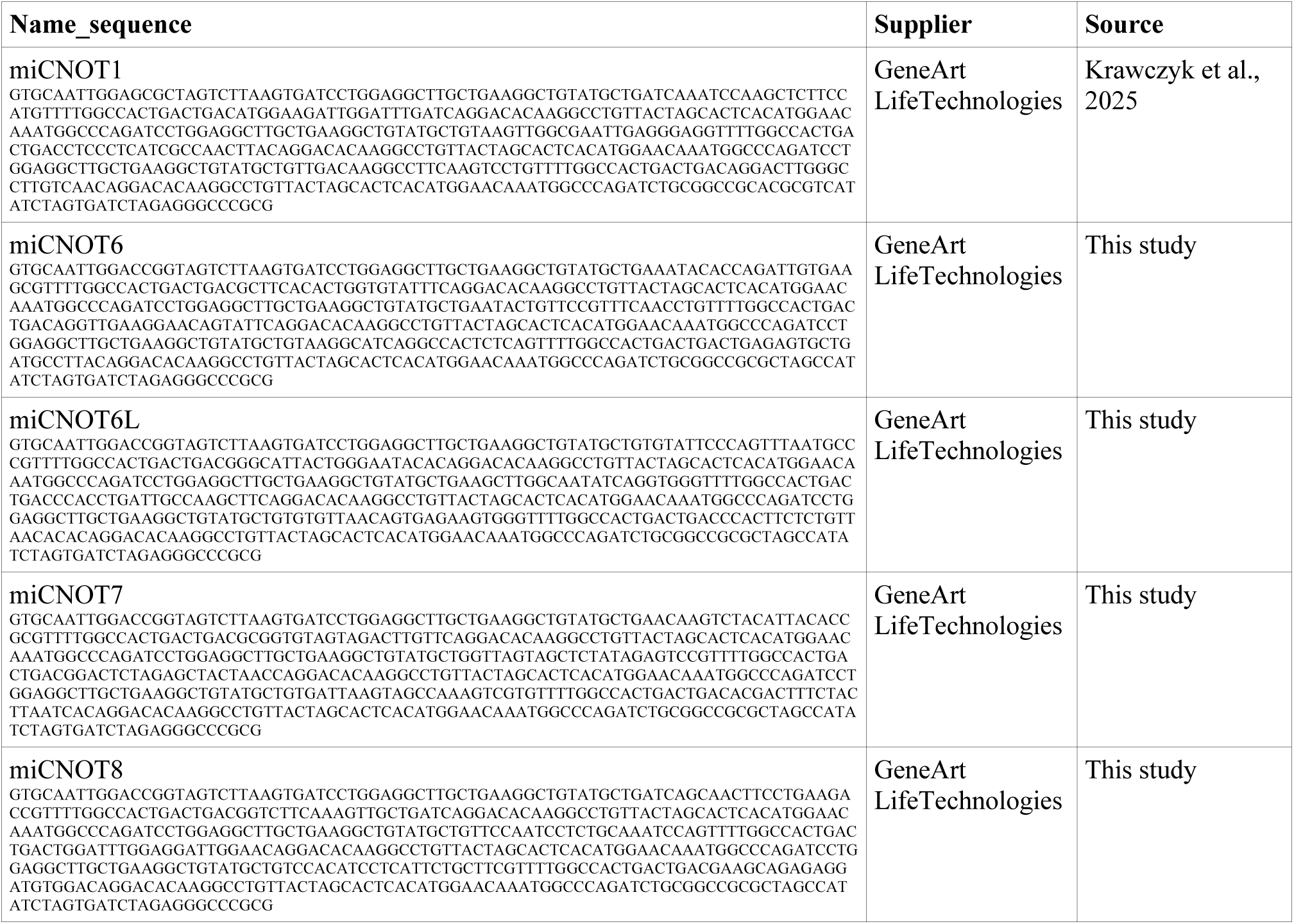
List of microRNA constructs used for silencing of CNOT1, CNOT6, CNOT6L, CNOT7 and CNOT8. Sequences are given as ordered from GeneArt service at Life Technologies. Inserts were digested using BshTI and NheI or BspT1 and NotI and inserted into pKK-B16 mCherry_EGFP vectors with bidirectional promoters (Szczęsny et al., 2018).

**Supplementary Table 3.** List of plasmids used in this work to construct stable HEK293 cell lines. All constructs were cloned into the pKK-B16 *BspT*I_*Not*I_mCherry/EGFP_*BshT*I_*Nhe*I plasmid, in which both cloning sites are flanking a bi-directional tetracycline-dependent promoter.

| **Plasmid name** | **Stable cell line** | **Source** |
|---|---|---|
| pKK-BI16_CNOT1miR:nucCHERRY_Promotor_EGFP-TEV-empty | HEK293 | Krawczyk et al., 2025 |
| pKK-BI16_CNOT6miR:nucCHERRY_Promotor_EGFP-TEV-empty | HEK293 | This study |
| pKK-BI16_CNOT6LmiR:nucCHERRY_Promotor_EGFP-TEV-empty | HEK293 | This study |
| pKK-BI16_CNOT6LmiR:nucCHERRY_Promotor_EGFP-TEV_miCNOT6 | HEK293 | This study |
| pKK-BI16_CNOT7miR:nucCHERRY_Promotor_EGFP-TEV-empty | HEK293 | This study |
| pKK-BI16_CNOT8miR:nucCHERRY_Promotor_EGFP-TEV-empty | HEK293 | This study |
| pKK-BI16_CNOT8miR:nucCHERRY_Promotor_EGFP-TEV_miCNOT7 | HEK293 | This study |

**Supplementary Table 4.**
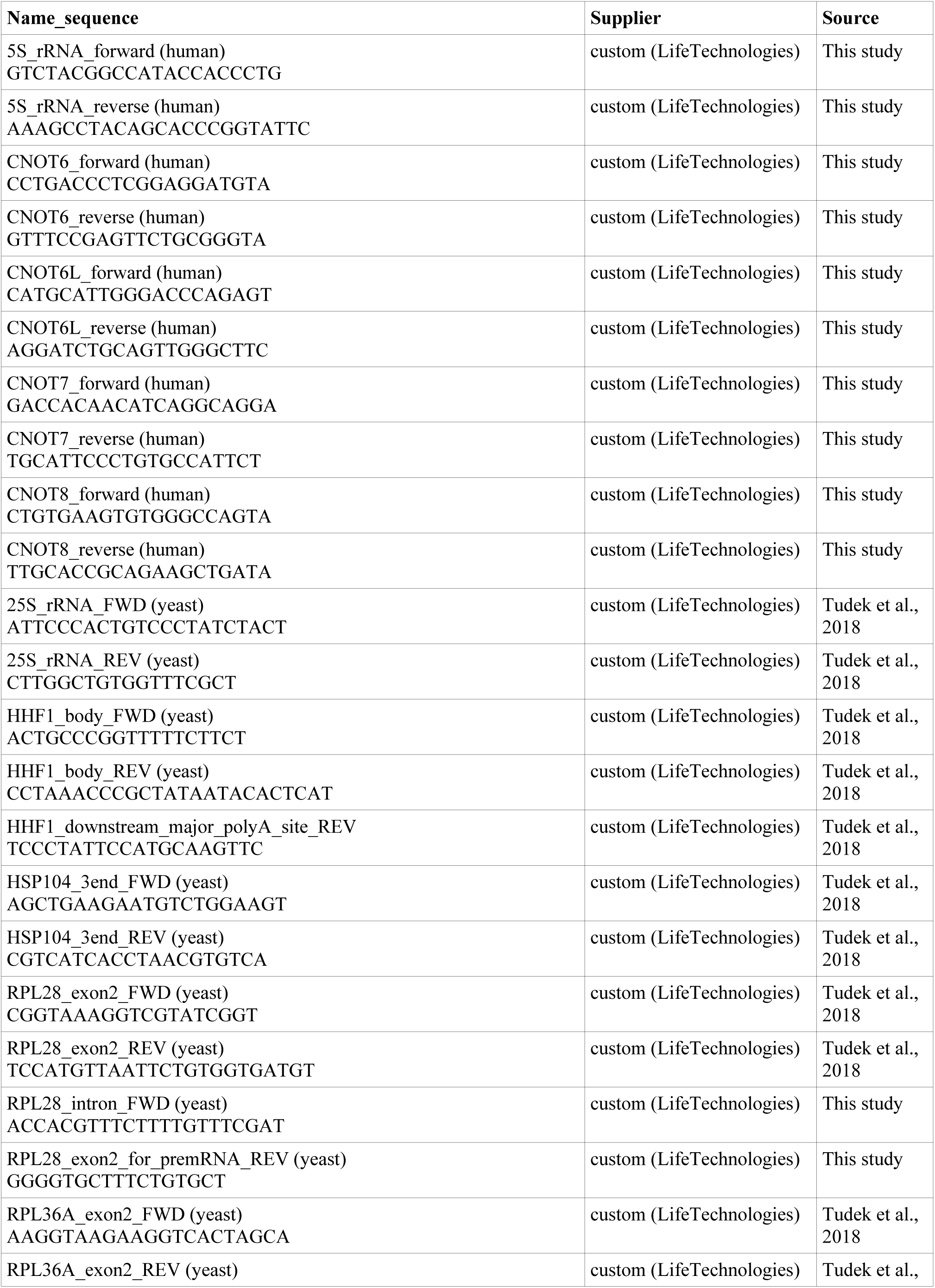
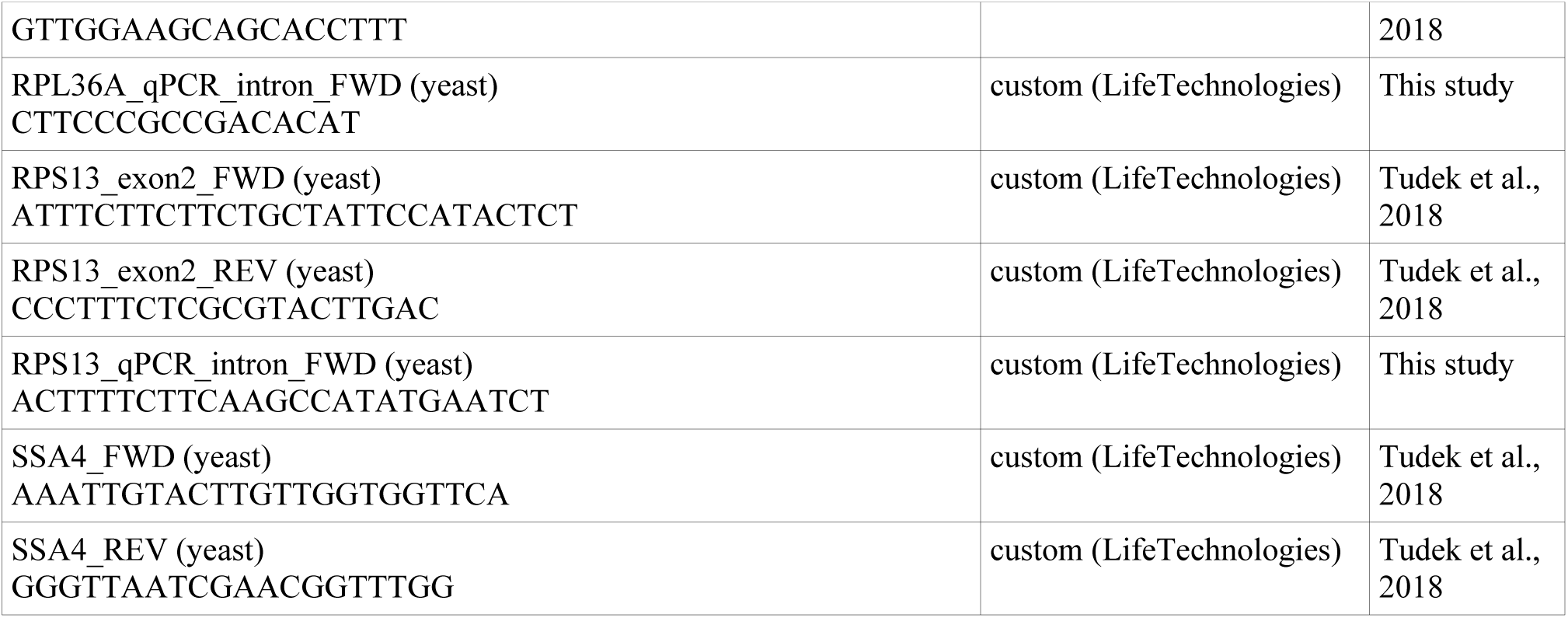
List of oligonucleotides used in this study. The list presents first the oligonucleotides annealing to human and then yeast targets.

**Supplementary Table 5.**
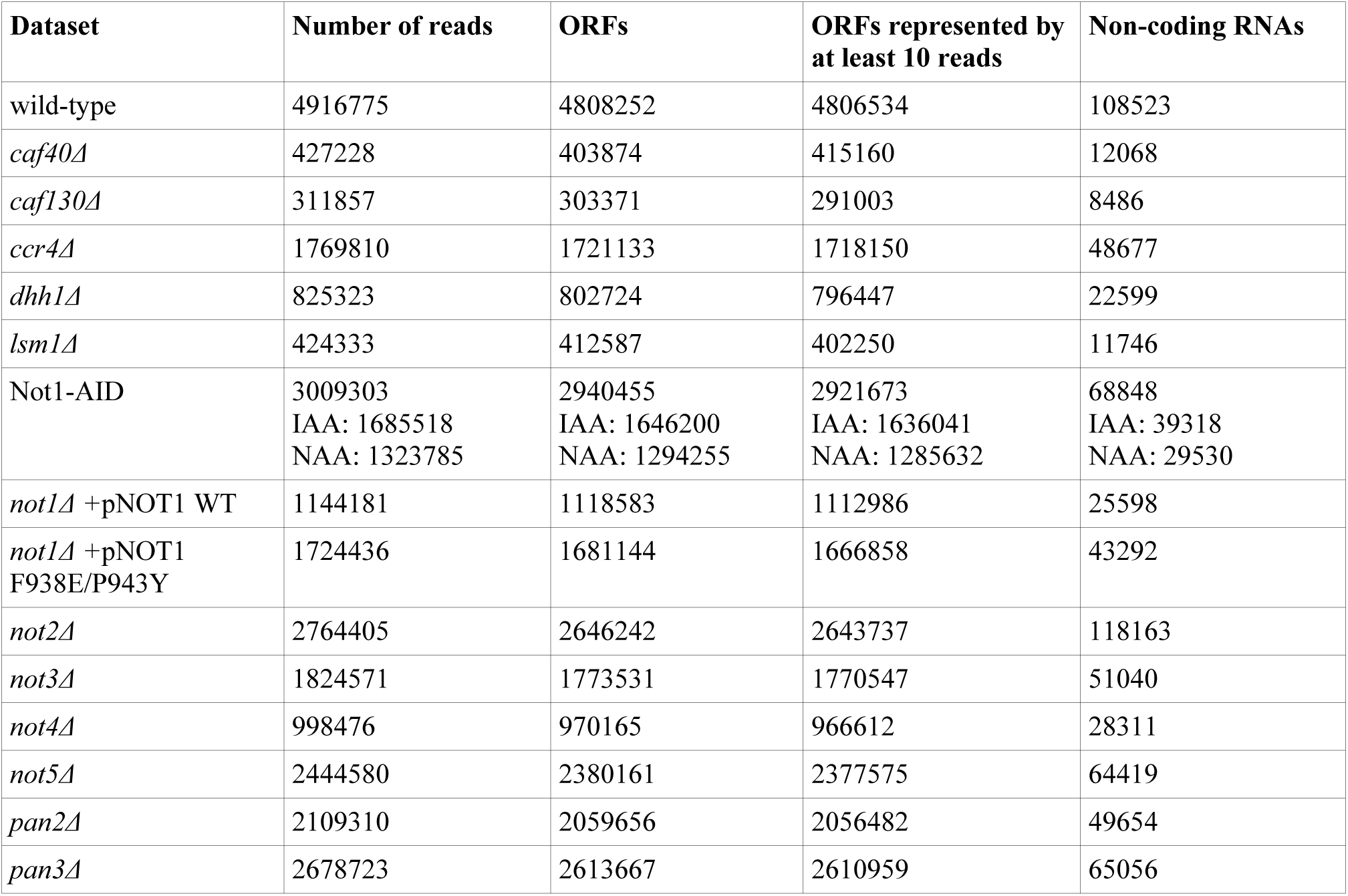
Basic statistics on the DRS datasets.

**Supplementary Table 6.**
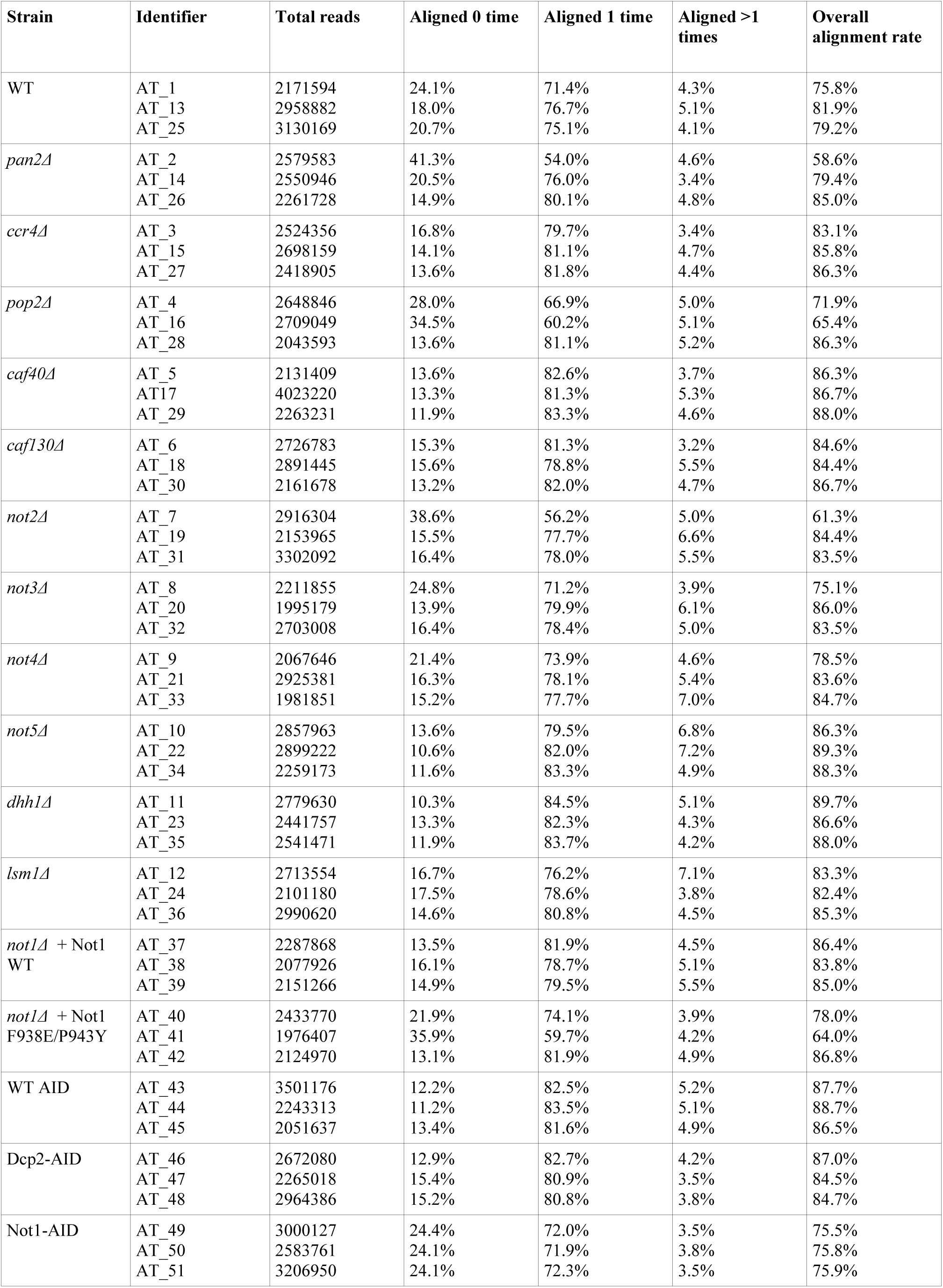
Basic statistics on the Illumina datasets.

| **Strain** | **Identifier** | **Total reads** | **Aligned 0 time** | **Aligned 1 time** | **Aligned >1 times** | **Overall alignment rate** |
|---|---|---|---|---|---|---|
| WT | AT_1 | 2171594 | 24.1% | 71.4% | 4.3% | 75.8% |
|  | AT_13 | 2958882 | 18.0% | 76.7% | 5.1% | 81.9% |
|  | AT_25 | 3130169 | 20.7% | 75.1% | 4.1% | 79.2% |
| *pan2Δ* | AT_2 | 2579583 | 41.3% | 54.0% | 4.6% | 58.6% |
|  | AT_14 | 2550946 | 20.5% | 76.0% | 3.4% | 79.4% |
|  | AT_26 | 2261728 | 14.9% | 80.1% | 4.8% | 85.0% |
| *ccr4Δ* | AT_3 | 2524356 | 16.8% | 79.7% | 3.4% | 83.1% |
|  | AT_15 | 2698159 | 14.1% | 81.1% | 4.7% | 85.8% |
|  | AT_27 | 2418905 | 13.6% | 81.8% | 4.4% | 86.3% |
| *pop2Δ* | AT_4 | 2648846 | 28.0% | 66.9% | 5.0% | 71.9% |
|  | AT_16 | 2709049 | 34.5% | 60.2% | 5.1% | 65.4% |
|  | AT_28 | 2043593 | 13.6% | 81.1% | 5.2% | 86.3% |
| *caf40Δ* | AT_5 | 2131409 | 13.6% | 82.6% | 3.7% | 86.3% |
|  | AT17 | 4023220 | 13.3% | 81.3% | 5.3% | 86.7% |
|  | AT_29 | 2263231 | 11.9% | 83.3% | 4.6% | 88.0% |
| *caf130Δ* | AT_6 | 2726783 | 15.3% | 81.3% | 3.2% | 84.6% |
|  | AT_18 | 2891445 | 15.6% | 78.8% | 5.5% | 84.4% |
|  | AT_30 | 2161678 | 13.2% | 82.0% | 4.7% | 86.7% |
| *not2Δ* | AT_7 | 2916304 | 38.6% | 56.2% | 5.0% | 61.3% |
|  | AT_19 | 2153965 | 15.5% | 77.7% | 6.6% | 84.4% |
|  | AT_31 | 3302092 | 16.4% | 78.0% | 5.5% | 83.5% |
| *not3Δ* | AT_8 | 2211855 | 24.8% | 71.2% | 3.9% | 75.1% |
|  | AT_20 | 1995179 | 13.9% | 79.9% | 6.1% | 86.0% |
|  | AT_32 | 2703008 | 16.4% | 78.4% | 5.0% | 83.5% |
| *not4Δ* | AT_9 | 2067646 | 21.4% | 73.9% | 4.6% | 78.5% |
|  | AT_21 | 2925381 | 16.3% | 78.1% | 5.4% | 83.6% |
|  | AT_33 | 1981851 | 15.2% | 77.7% | 7.0% | 84.7% |
| *not5Δ* | AT_10 | 2857963 | 13.6% | 79.5% | 6.8% | 86.3% |
|  | AT_22 | 2899222 | 10.6% | 82.0% | 7.2% | 89.3% |
|  | AT_34 | 2259173 | 11.6% | 83.3% | 4.9% | 88.3% |
| *dhh1Δ* | AT_11 | 2779630 | 10.3% | 84.5% | 5.1% | 89.7% |
|  | AT_23 | 2441757 | 13.3% | 82.3% | 4.3% | 86.6% |
|  | AT_35 | 2541471 | 11.9% | 83.7% | 4.2% | 88.0% |
| *lsm1Δ* | AT_12 | 2713554 | 16.7% | 76.2% | 7.1% | 83.3% |
|  | AT_24 | 2101180 | 17.5% | 78.6% | 3.8% | 82.4% |
|  | AT_36 | 2990620 | 14.6% | 80.8% | 4.5% | 85.3% |
| *not1Δ* + Not1 WT | AT_37 | 2287868 | 13.5% | 81.9% | 4.5% | 86.4% |
|  | AT_38 | 2077926 | 16.1% | 78.7% | 5.1% | 83.8% |
|  | AT_39 | 2151266 | 14.9% | 79.5% | 5.5% | 85.0% |
| *not1Δ* + Not1 F938E/P943Y | AT_40 | 2433770 | 21.9% | 74.1% | 3.9% | 78.0% |
|  | AT_41 | 1976407 | 35.9% | 59.7% | 4.2% | 64.0% |
|  | AT_42 | 2124970 | 13.1% | 81.9% | 4.9% | 86.8% |
| WT AID | AT_43 | 3501176 | 12.2% | 82.5% | 5.2% | 87.7% |
|  | AT_44 | 2243313 | 11.2% | 83.5% | 5.1% | 88.7% |
|  | AT_45 | 2051637 | 13.4% | 81.6% | 4.9% | 86.5% |
| Dcp2-AID | AT_46 | 2672080 | 12.9% | 82.7% | 4.2% | 87.0% |
|  | AT_47 | 2265018 | 15.4% | 80.9% | 3.5% | 84.5% |
|  | AT_48 | 2964386 | 15.2% | 80.8% | 3.8% | 84.7% |
| Not1-AID | AT_49 | 3000127 | 24.4% | 72.0% | 3.5% | 75.5% |
|  | AT_50 | 2583761 | 24.1% | 71.9% | 3.8% | 75.8% |
|  | AT_51 | 3206950 | 24.1% | 72.3% | 3.5% | 75.9% |

